# Aggregated A-synuclein leads to corticostriatal synaptic dysfunction

**DOI:** 10.1101/2025.01.29.635597

**Authors:** Charlotte F. Brzozowski, Harshita Challa, Nolwazi Z. Gcwensa, Dominic Hall, Douglas Nabert, Nicole Chambers, Ignacio Gallardo, Michael Millet, Laura Volpicelli-Daley, Mark S. Moehle

## Abstract

Neuronal inclusions of α-synuclein (α-syn) are pathological hallmarks of Parkinson’s disease (PD) and Dementia with Lewy Bodies (DLB). α-Syn pathology accumulates in cortical neurons which project to the striatum. To begin to understand how α-syn pathology effects cortico-striatal synapses, pre-formed α-syn fibrils (PFF) were injected into the striatum to induce robust α-syn aggregation in corticostriatal-projecting neurons. Electrophysiological recordings of striatal spiny projection neurons (SPNs) acute slices found a significant decrease in evoked corticostriatal glutamate release in mice with PFF-induced aggregates compared to monomer injected mice. Expansion microscopy, confocal microscopy and Imaris reconstructions were used to identify vGLUT1 positive presynaptic terminals juxtaposed to Homer-positive postsynaptic densities, termed synaptic foci. Quantitation of synaptic loci density revealed a loss of corticostriatal synapses. Immunoblots of the striatum show reductions in expression of pre-synaptic proteins with selective reduction in AMPA and NMDA receptor subunits in mice with α-syn aggregates compared to controls. Paradoxically, a small percentage of remaining VLGUT1+ synaptic loci with small, intrasynaptic α-syn aggregates showed enlarged volumes compared to nearby synapses without α-syn aggregates. Our combined physiology and high-resolution imaging data point to dysfunction of corticostriatal synapses in mice harboring □-synuclein inclusions, which may contribute to impaired basal ganglia circuitry in PD.

**Highlights:**

- Corticostriatal glutamate drive is impaired in the presence of pathological α-syn
- α-Syn aggregation causes early loss of corticostriatal synapses
- Synaptic loci positive for small α-syn aggregates show volume increases
- Striatal expression of select synaptic proteins are reduced in animals with α-syn pathology

## Introduction

□-Synuclein (□-syn) primarily localizes to presynaptic terminals and is particularly enriched in glutamatergic excitatory neurons (Brzozowski et al., 2021; Taguchi et al., 2016). In these synapses‘, □-syn interacts with the SNARE complex to modulate synaptic transmission and excitatory neurotransmitter vesicular dynamics, suggesting a unique role for □-syn in modulating glutamate release (Burré et al., 2010; Sharma and Burré, 2023; Sun et al., 2019). However, in Parkinson’s Disease (PD) and Dementia with Lewy Bodies (DLB), □-syn forms hallmark pathological aggregates called Lewy Bodies and Neurites (together Lewy Pathology, LP) (Henderson et al., 2019; Outeiro et al., 2019; Spillantini et al., 1997). Despite the clear pathological link between □-syn and PD, the impact of LP on synaptic physiology is poorly understood.

In pathological states, □-syn forms into toxic amyloid fibrils (Firbank et al., 2016; Froula et al., 2019; Yang et al., 2022), which have been shown to localize to synaptic terminals in the cortex of postmortem DLB brain (Colom-Cadena et al., 2017; Schulz-Schaeffer, 2010). Studies using templated □-syn aggregation models and post-mortem tissue from PD and DLB patients show a partial loss of native □-syn at the presynaptic terminal (Chen et al., 2022) and show reduced expression of SNARE proteins VAMP2 and SNAP25 in excitatory layer 5 pyramidal neurons and cultured neurons (Goralski et al., 2024; Volpicelli-Daley et al., 2011). Post-synaptically, there is decreased number and remodeled ultrastructure of dendritic spines (Blumenstock et al., 2017; Froula et al., 2018; Villalba and Smith, 2018). These findings have suggested a unique role for □-syn aggregates decreasing synaptic architecture and function, but this has been largely unexplored in pathologically relevant models.

To begin to address this question, we utilized the preformed fibril (PFF) mouse model to investigate alterations in corticostriatal synapses at an early timepoint after formation of α-syn aggregates. Using a combination of physiological recordings and high-resolution imaging with 3D reconstruction of synaptic loci, we found early defects in corticostriatal synaptic transmission and loss of cortical-striatal synapse density in the striatum upon α-syn pathology formation. Paradoxically, remaining presynaptic terminals containing α-synuclein aggregates showed increased volumes. Additionally, we found reduced expression levels of pre-synaptic markers and selective reduction in NMDA receptor subunits in the striatum of PFF-injected animals which may underlie functional changes in corticostriatal transmission observed. Together, our data suggest decreased cortico-striatal function that rapidly develops upon □-syn aggregation and further suggests that cortico-striatal synaptic efficacy may be early drivers of symptoms in PD and DLB.

## Methods and Materials

### Animals

Wildtype C57BL/6J mice were acquired from the Jackson Laboratory (strain 000664). The mice followed a 12-hour light/dark cycle with unrestricted access to food and water according to NIH guidelines for care and use of research animals. Animal protocols were approved by the respective Institutional Animal Care and Use Committees at the University of Alabama at Birmingham and the University of Florida.

### Preparation of α-syn preformed fibrils (PFF)

Mouse monomeric α-syn was purified in E. coli as described in Volpicelli-Daley et al., 2014(Volpicelli-Daley et al., 2014). Endotoxin removal was facilitated using Pierce High-Capacity Endotoxin RemovalSpin columns and endotoxin levels (<10 EU/mg) were subsequently determined using the LAL endotoxin assay kit (GeneScript,). Protein concentration was measured at 280 absorbance using an extinction factor (□) 7450 M^−1^ cm^−1^ for mouse monomeric α-syn. To generate PFFs, 300 μM monomeric α-syn underwent constant shaking at 37 °C for 7 days in a potassium chloride/Tris buffer (150 mM KCl, 50 mM Tris-HCl, pH 7.5). PFFs were isolated from soluble protein through centrifugation at 13200 rpm for 10 minutes at room temperature. PFF concentration was measured after dissociation in guanidinium chloride solution (8M) into monomeric α-syn, and protein concentration measured as mentioned above. Thereafter, PFFs underwent resuspension to a final concentration of 300 μM protein in 150 mM KCl and 50 mM Tris-HCl buffer. Fragmentation of PFFs into smaller, seeding-competent fragments was achieved by sonication using a Qsonica 700W cup horn sonicator, for 15 minutes at 30% amplitude, employing 3-second on and 2-second off pulses. The fragmentation of PFFs to a desired size between 20 nm and 80 nm was confirmed through dynamic light scattering measurements (Dynapro Nanostar, Wyatt Technologies) for quality control. Fragmented fibrils were subsequently used in experiments. All experiments involving PFFs followed rigorous safety guidelines and decontamination procedures (Bousset et al., 2016).

### Stereotaxic surgeries

Surgeries and intracranial injections of PFFs (or control) were performed on 3–4-month-old C57BL/6J mice using a digital stereotaxic frame. Mice were deeply anesthetized with isoflurane and mouse respiration was monitored throughout the procedure. Controlled by a digital pump, a gas-tight, 26s-gauge needle (Hamilton) was used to inject PFFs, or either control monomeric □-syn or control PBS vehicle solution. A total of 10 μg protein at 5mg/mL concentration per hemisphere was injected into the striatum (Physiology experiments at UF, coordinates: ML: +2.0; AP: +0.2, DV: −3.0 from bregma; molecular biology experiments at UAB, coordinates: ML: +2.0, AP: +1.0 from bregma, DV: −3.2 from dura) at 0.5 µL/min. The needle was held in place for 5 minutes after the injection ended then slowly withdrawn.

### Immunofluorescence

#### Transcardial perfusions

C57BL/6J mice were deeply anesthetized with isoflurane and transcardially perfused with ice-cold 0.9% saline, containing sodium nitroprusside (0.5% w/v) and 10 units/mL heparin. Brains were removed from skull and hemi-dissected on ice. The hemispheres designated for immunofluorescence and expansion microscopy experiments underwent fixation in 4% paraformaldehyde (PFA) in phosphate-buffered saline (PBS) for 48h at 4 °C followed by immersion 30% sucrose in PBS followed by snap-freezing and storage at −80°C. Brains were sectioned at 40 μM thickness on a freezing microtome (Leica SM 2010R), and stored in a cryoprotectant solution at −20 °C.

#### Immunofluorescence of Brain Sections

Sections were rinsed six times in tris-buffered saline (TBS) followed by an antigen retrieval step (10 mM sodium citrate, 0.05% Tween-20, pH 6.0) for 1 hour at 37 °C. Brain sections were blocked and permeabilized for 1 hour at 4 °C in 5% goat serum and 0.1% TritonX-100 in TBS. Both primary and secondary antibodies (Table 1 and 2) were diluted in 5% goat serum in TBS. Sections were incubated in primary antibody solution overnight at 4 °C with shaking. After three rinses in TBS, sections were incubated in Alexa-Fluor conjugated secondary antibodies (Table X) for 2 hours at 4 °C followed by mounting onto glass slides (Superfrost Plus) using Prolong Gold.

**Table 1:** Overview of Primary Antibodies.

| Antibody Target | Supplier | Catalog No. | RRID | Host Species/ Isotype/ Mono or Polyclonal | Method (Dilution) |
| --- | --- | --- | --- | --- | --- |
| DARPP-32 | R&D | MAB4230 | RRID:AB_2169021 | Monoclonal Rat IgG2a | IF (1:500) |
| DAT | Milipore | MAB369 | RRID:AB_2190413 | Monoclonal Rat IgG2a kappa | IF (1:1000) |
| GAPDH | abcam | ab181603 | RRID:AB_2687666 | Monoclonal Rabbit IgG | IB (1:10000) |
| GluA1 | abcam | ab109450 | RRID:AB_10860361 | Monoclonal Rabbit IgG | IB (1:2000) |
| GluA2 | abcam | ab133477 | RRID:AB_2620181 | Monoclonal Rabbit IgG | IB (1:1000) |
| GluA3 | Synaptic Systems | 182204 | RRID:AB_2247891 | Polyclonal Rabbit | IB (1:1000) |
| Homer1 | Synaptic Systems | 160006 | RRID:AB_2631222 | Polyclonal Chicken | IB (1:2000)<br>IF (1:1000)<br>ExM (1:500) |
| NeuN | Millipore | ABN91 | RRID:AB_11205760 | Polyclonal Chicken IgY | IF (1:2500) |
| NMDAR1 | abcam | ab109182 | RRID:AB_10862307 | Monoclonal Rabbit IgG | IB (1:5000) |
| NMDAR2a | abcam | ab133265 | RRID:AB_11158532 | Monoclonal Rabbit IgG | IB (1:1000) |
| NMDAR2b | abcam | ab254356 | RRID:AB_2895296 | Monoclonal Rabbit IgG | IB (1:2000) |
| pS129-a-syn | abcam | ab51253 | RRID:AB_869973 | Monoclonal Rabbit IgG | IF (1:5000)<br>ExM (1:2500) |
| Snap25 | Sigma | S9684 | RRID:AB_261576 | Polyclonal Rabbit IgG | IB (1:1000) |
| TH | Millipore | AB152 | RRID:AB_390204 | Polyclonal Rabbit IgG | IB (1:2000)<br>IF (1:2000) |
| tubulin | Santa Cruz | SC23947 | RRID:AB_628410 | Monoclonal Mouse IgG1 | IB (1:2000) |
| VAMP2 | Synaptic Systems | 104211 | RRID:AB_887811 | Monoclonal Mouse IgG1 | IB (1:1000) |
| VGLUT1 | Synaptic Systems | 135304 | RRID:AB_887878 | Polyclonal Guinea Pig antiserum | IB (1:2000)<br>IF (1:1000)<br>ExM (1:500) |
| VGLUT2 | Synaptic Systems | 135421 | RRID:AB_2619823 | Monoclonal Mouse IgG2a | IB (1:2000)<br>IF (1:1000) |
| Vinculin | Bio-Rad | MCA465GA | RRID:AB_3100992 | Monoclonal Mouse IgG1 | IB is (1:1000) |

**Table 2:** Overview of Secondary Antibodies.

| Fluorophore/Enzyme | Target Species/ | Supplier | Catalog No. | RRID | Method (Dilution) |
| --- | --- | --- | --- | --- | --- |
|  | Isotype |  |  |  |  |
| Alexa Fluor™ 488 | chicken IgY | ThermoFisher | A-11039 | RRID:AB_2534096 | ExM (1:250)<br>IF (1:500) |
| Alexa Fluor™ 488 | rabbit IgG | ThermoFisher | A-11034 | RRID:AB_2576217 | IF (1:500) |
| Alexa Fluor™ 555 | guinea pig | ThermoFisher | A-21435 | RRID:AB_2535856 | ExM (1:250)<br>IF (1:500) |
| Alexa Fluor™ 555 | rat IgG | ThermoFisher | A-21434 | RRID:AB_2535855 | IF (1:500) |
| Alexa Fluor™ 568 | mouse IgG2a | ThermoFisher | A-21134 | RRID:AB_2535773 | IF (1:500) |
| Alexa Fluor™ 647 | chicken | ThermoFisher | A-21449 | RRID:AB_2535866 | IF (1:500) |
| Alexa Fluor™ 647 | rabbit IgG | ThermoFisher | A-21245 | RRID:AB_2535813 | ExM (1:250)<br>IF (1:500) |
| Alexa Fluor™ 680 | mouse IgG | ThermoFisher | A-21058 | RRID:AB_2535724 | IB (1:5000) |
| Atto 647N | rabbit | Sigma | 40839 | RRID:AB_1137669 | IB (1:5000) |
| HRP | chicken | ThermoFisher | SA1-300 | RRID:AB_325995 | IB (1:5000) |
| HRP | mouse IgG | ThermoFisher | SA1-100 | RRID:AB_325993 | IB (1:5000) |
| HRP | rabbit | ThermoFisher | A-16023 | RRID:AB_2534697 | IB (1:5000) |
| IRDye 680RD | guinea pig | LI-COR | 925-68077 | RRID:AB_2814914 | IB (1:5000) |
| IRDye 800CW | rabbit | LI-COR | 926-32211 | RRID:AB_621843 | IB (1:5000) |

#### Expansion Microscopy of Brain sections

Brain sections were prepared for immunofluorescence as described above. After the secondary antibody incubation step and washes, pre-expansion imaging using a wide field fluorescent microscope was performed by imaging whole hemisphere signal for DAPI. ExM protocol was followed as described in Bucur et al 2020 (Bucur et al., 2020). Sections were then anchored in 0.1 TritonX-100/acryloyl X (Life Tech Corp) solution for 4 hours at room temperature, followed by embedding in a swellable biopolymer network consisting of monomer solution (8.6g/100mL sodium acrylate, 2.5 g/100 mL acrylamide, 0.1 g/100mL N, N-methylenebisacrylamide, 11.7 g/100 mL NaCl, in 1XPBS) with the addition of 0.2% (w/v) APS, 0.2% (w/v) TEMED, and 0.01% (w/v) 4-Hydroxy tempo. Post polymerization, gels underwent proteinase K treatment (NEB, 1:200 dilution) in digestion buffer (60 mM Tris, 25 mM EDTA, 0.5% v/v Triton X-100, 0.8 M NaCl, pH 8.0) for 45 minutes at 60°C to allow for isotropic expansion followed by five washes in 1X PBS to allow for expansion of the tissue. The expanded tissue sections were re-imaged using a widefield fluorescence microscope.

For calculating the expansion factor of expanded samples, DAPI-stained, whole hemispheres of pre-expansion samples and post-expansion samples were imaged using a 10X air objective with 5% overlap between stitched images for tiling.

### Fluorescent Microscopy and Image analysis

#### Wide Field Fluorescence Microscopy

Whole hemisphere tiled images for visualization of pS129-α-syn-positive aggregates were generated using an inverted ZEISS Axiovert.Z1 Cell Observer microscope with a Hamamatsu Orca Flash high-speed camera. Single images obtained with a 20X air objective (LD Plan Neofluoar 20X/0.4 Corr M27) were stitched using Carl Zeiss software with 5% overlap between image tiles. Single frame images of pS129-positive aggregates were acquired with a 20X (low magnification, Zeiss LD Plan Neofluoar 20X/0.4 Corr M27) and 40X (high magnification, Zeiss LD plan Neofluoar 40X/0.6 Ph2 Corr) air objectives. Image brightness and contrast were adjusted using Carl Zeiss software look-up table (LUT) settings.

#### Confocal Microscopy

An inverted Nikon A1R confocal microscope was used for all confocal imaging. *Synapse analysis using non-expanded tissue:* Non-expanded brain sections were imaged using a 60X oil immersion objective (Nikon CFI Plan Apochromat λD 60X/1.42). Ten images frames were collected for at least 2 separate coronal slices in the dorsal striatum of each animal (2 µm z-stacks, step size 0.125 µm) using Nyquist settings (1024 x 1024 resolution, 2.06 optical zoom) To prevent imaging in areas close to injection needle tract, striatal sections with a minimal distance of 100 µm to injection site (AP 0.9-0.00 from bregma) were selected for imaging. *Synapse analysis using expanded tissue:* Expanded brain sections were imaged with an apochromatic, long working distance 40X water immersion objective (Nikon Apo LWD 40X/1.15 WI). 3-5 image frames were collected in the dorsal striatum of each animal (4 µm z-stacks, step size 0.125 µm). Images were collected acquired using Nyquist settings (1024 x 1024 resolution, 2.06 optical zoom) for all channels. Images determined to have a significant extent of signal drift due to sample shifting during data collection were omitted from further analysis. Thereafter, z-stacks were deconvolved with NIS Advanced Software (Nikon) using the Lucy-Richardson deconvolution algorithm. Laser intensity and detector gain were kept constant throughout experimental sets for imaging of both expanded and non-expanded tissue.

*Tyrosine hydroxylase (TH) fiber density:* TH and NeuN fluorescent signal in non-expanded tissue sections was determined using a 20X air objective. Four Single-plane image frames were collected in the dorsal striatum of 4 separate coronal sections (AP –0.2 −0.0 from bregma) in each animal. Images were acquired using 2024 x 2024 resolution with no optical zoom. Laser intensity and detector gain were kept constant throughout the experiment.

#### Image Analysis

Synapse analysis: Confocal z-stacks were deconvolved (Lucy Richardson, 20 iterations) using advanced NIS analysis software (Gcwensa et al., 2024). Deconvolved z-stacks were converted into IMS files using Imaris software V.9 (Andor technologies) and further processed in Imaris for surface reconstruction. In short, surfaces were reconstructed from thresholded, fluorescent signal and synaptic loci defined as pre and postsynaptic surfaces with the shortest distance to nearest neighboring surface filter set below 0.01 µm. Total and synaptic surface densities were calculated by dividing surface count by total z-stack volume followed by averaging frame values for each animal. Total z-stack volumes were computed by generating a surface filling the entire z-stack and using the volume provided by Imaris software. For morphology parameters, mean values for each imaging frame were averaged, per animal. All Imaris surface construction and analysis parameters were kept constant within experiments.

### Immunoblot analysis

#### Protein lysate generation

Anesthetized C57BL/6J mice were transcardially perfused with ice-cold 0.9% saline, containing sodium nitroprusside (0.5% w/v) and 10 units/mL heparin. Thereafter, brains were hemi-dissected on ice, and the right hemisphere was further processed for immunofluorescence procedures. Left hemispheres of bilaterally injected brains were further dissected on ice into midbrain, hippocampus, striatum and cortex and flash frozen in a dry ice/EtOH slurry. Striatal lysates were generated by homogenization in 1% Triton-X/1XTBS solution containing protease (Thermo Scientific Pierce Protease Inhibitor Mini Tablets) and phosphatase inhibitors (Thermo Scientific Pierce Phosphatase Inhibitor Mini Tablets). Triton-soluble and Triton-insoluble fractions were isolated as described in Volpicelli et al. 2011. In short, after homogenization of samples in 1% Triton-X/1XTBS solution, samples were centrifuged at maximal speed (15000 rpm) for 30 minutes at 4 °C and supernatant was collected as Triton-X soluble fraction. Protein concentration in lysates was quantified using a BCA assay. Samples were diluted to a 5 mg/mL protein concentration in laemmli buffer/10% Dithiothreitol. Striatal protein lysates underwent electrophoresis using precast, 4-20% gradient SDS-PAGE gels and transferred overnight onto PVDF membranes. Membranes were blocked in Everyblot blocking buffer and incubated in primary antibody (table 1) solutions in either Everyblot solution or 2% BSA/0.2% Tween in TBS. Membranes were incubated in HRP-coupled secondary antibodies (table 2) and developed using ECL solution. VGLUT1 and VGLUT2 blots and respective loading controls were incubated in infrared, fluorophore-coupled secondary antibodies for 2 hours. All blots were imaged in a Biorad imaging device using either chemiluminescence imaging (for HRP-conjugated secondary detection) or infrared, fluorescent imaging in the 680 and 800 wavelength channels for detection of fluorophore-conjugated secondary antibodies. The band intensities for proteins of interest and respective loading control protein bands were measured by using the plot band function in ImageJ followed by normalization to respective loading control.

### Ex vivo Electrophysiology

#### Preparation of slices

Acute slices were prepared as described in Chambers et al. 2024 (Chambers et al., 2024). In short, mice were deeply anesthetized with isoflurane, and then perfused with ice cold, 95% O_2_ and 5% CO_2_ gas-equilibrated N-Methyl-D-gluconate (93 mM NMDG, 2.5 mM KCl, 1.4 mM Na-Phosphate Monobasic Monohydrate, 30 mM Na-bicarbonate, 20 mM Hepes, 5 mM Glucose, 2 mM thiourea, 3 mM Na-Pyruvate, 10 mM MgSO_4_, 0.5 mM CaCl_2_) solution. Brains were harvested and 250 µm coronal brain slices containing the striatum were generated using a vibratome (VT1200S, Leica) while sections remained immersed in ice cold, oxygenated recovery solution. Sections were placed in warm, equilibrated NMDG-solution (33 °C) for 10 minutes and then transferred into a sodium ascorbate (5mM) and artificial cerebrospinal fluid (aCSF: (126 mM NaCl, 2.5 mM KCl, 1.4 mM Na-Phosphate Monobasic Monohydrate, 26 mM Na-bicarbonate, 1 mM Glucose, 1.5 mM MgSO_4_, 2 mM CaCl_2_) equilibrated with 95% O_2_ and 5% CO_2_ solution for recovery at RT for at least an hour until use.

#### Electrophysiological recordings

Coronal sections were transferred into the recording chamber and perfused at 5mL/min with warm aCSF (95% O_2_/ 5% CO_2_ saturated) using feedback controlled in-line heating element. Neurons were visualized using IR-DIC Optics with a DAGE-2000 infrared camera. Whole cell patch clamp recordings of SPNs in the dorsal striatum were performed using glass borosilicate pipettes (3-8mOhm resistance) filled with internal K-gluconate solution (125 mM K-gluconate, NaCl 4mM, 10 mM HEPES, 4 mM Mg-ATP, 0.3 mM Na-ATP, 10 mM Tris-phosphate). The inclusion criteria to identify SPNs were a resting membrane potential around −80mV, absence of spontaneous action potential (AP) firing, and non-adapting AP firing frequency during depolarization. For spontaneous excitatory transmission experiments, neurons were held at −75 mV in current clamp configuration, and currents were recorded for 5 minutes. Neurons were then depolarized in 50 pA increments (1s of stimulation) in voltage clamp configuration up to 700 pA. Corticostriatal glutamate transmission experiments were performed by placing a concentric bipolar electrode connected to a stimulation unit on the corpus callosum (CC) white matter tract. Evoked excitatory postsynaptic currents (EPSC) in SPNs (holding potential −75 mV) from CC stimulation were measured in current clamp configuration. Rheobase current was assessed in voltage clamp configuration. Passive membrane properties were recorded using membrane test. Acute brain slices from physiological recordings were immediately postfixed in 4% PFA in PBS for 30 minutes at room temperature.

#### Electrophysiology data analysis

sEPSC: Current traces from 5-minute increment recordings were analyzed for spontaneous excitatory transmission using Easy Electrophysiology software. To detect sEPSCs, 3 different EPSC templates and a −15pA threshold respectively to baseline signal were used to semi-automatically detect sEPSCs. Detected event amplitude and inter event interval (IEI) were used to assess overall excitatory synaptic activity. The same templates were used for all experimental conditions. Evoked EPSC: Data was analyzed using Clampfit Software. First, baseline was subtracted from all traces and amplitudes of all EPSCs were measured using the event detection tool. Action potential (AP) analysis was performed using the event detection tool in Clampfit Software. For AP phase plots analysis, the average AP shape of evoked APs during 1ms 350 pA depolarization step were generated and phase plots created by plotting the time derivative of voltage change (dV/dt) over membrane voltage.

### Statistics

Statistical data analysis and graphs were generated using GraphPad Prism software. If not otherwise indicated in figure legends, all data is represented as mean□SEM. Prior to running statistical analysis, all data was subject to outlier testing using the Grout outlier test. Assumptions of respective statistical tests were assessed prior to running tests and are described in respective paragraphs. All statistical tests and analysis, mean□SEM and outliers are reported in statistical summary tables corresponding to each figure. <u>Unpaired Student t-test</u>: For two-column statistical analysis, two-tailed unpaired t-tests were used to assess statistical differences between groups (alpha set to 0.05). Normality was assessed using Shapiro-Wilk test and non-normally distributed data was analyzed using Mann-Whitney (MW) test. F-test was conducted to assess equality of variances between groups and a Welch t-test was used for data with statistically unequal variances. <u>One-way ANOVA:</u> To assess for a statistical main difference among means of three or more groups, one-way ANOVA test was used. Normal distribution was assessed using Shapiro-Wilk test, and non-normally distributed data was analyzed using Kruskal-Wallis nonparametric test. Equality of variance was assessed by Brown-Forsythe test, and if variances were found significantly different, Brown-Forsythe or Welch correction was applied. For significant one-way ANOVA results, appropriate post-hoc statistical testing was used to test for specific statistical differences between groups (Tukey post hoc test for ordinary one-way ANOVA, Dunnett post hoc test for Kruskal Wallis, indicated in respective statistical summary tables). <u>Two-way ANOVA:</u> This test was used to test for row and column main effect and row/column interaction in grouped analysis with two factors. Heteroscedacity of residuals was assessed using Spearman correlation coefficient and gaussian distribution of residuals was assessed using Shapiro-Wilk test. For positive 2-way-ANOVA interaction, Tukey multiple comparison post hoc test was used to assess statistical differences of the means in the different groups. <u>Repeated</u> <u>measure ANOVA:</u> For data analysis with repeated measure design, either a 2-way RM-ANOVA or mixed effects model analysis was used, while Geisser-Greenhouse correction was applied. For significant mixed effect model results, Šidák’s multiple comparison post hoc test was used to assess the significant differences between treatment groups of the repeated measure design.

## Results

### pS129-α-syn-positive aggregates accumulate in corticostriatal circuitry

To detect aggregated α-syn, an antibody that recognizes α-syn phosphorylated at serine 129 was used. Six weeks after intrastriatal injections of PFFs of pS129-α-syn-positive aggregates were visible in the motor cortices (primary motor cortex MOp, secondary motor cortex MOs), primary somatosensory area (SSp), anterior cingulate area and piriform area (Fig 1a, b). Injection of monomer into the striatum did not produce pS129-α-syn-positive aggregates (Fig S1). Somal and neuritic inclusions were enriched in layer V neurons (Fig1c). In the striatum, the majority of pathology appeared neuritic while some cells harbored somal pS129-α-syn in the dorsal striatum (Fig1d). Co-staining of SPN marker DARPP-32 and pS129-α-syn in the dorsal striatum showed somal aggregate formation in SPNs 6-weeks post-PFF injections (Fig S2a). Axonal projections to the striatum were positive for pS129-α-syn pathology, as shown by co-staining of pS129-α-syn and axonal marker Neurofilament (NF) to visualize axon collaterals in the CC (Fig S2b). Overall, these data showed that intrastriatal injections of PFFs produced neuritic and somal α-syn aggregates in layer V cortex and striatum.

**Figure 1:**
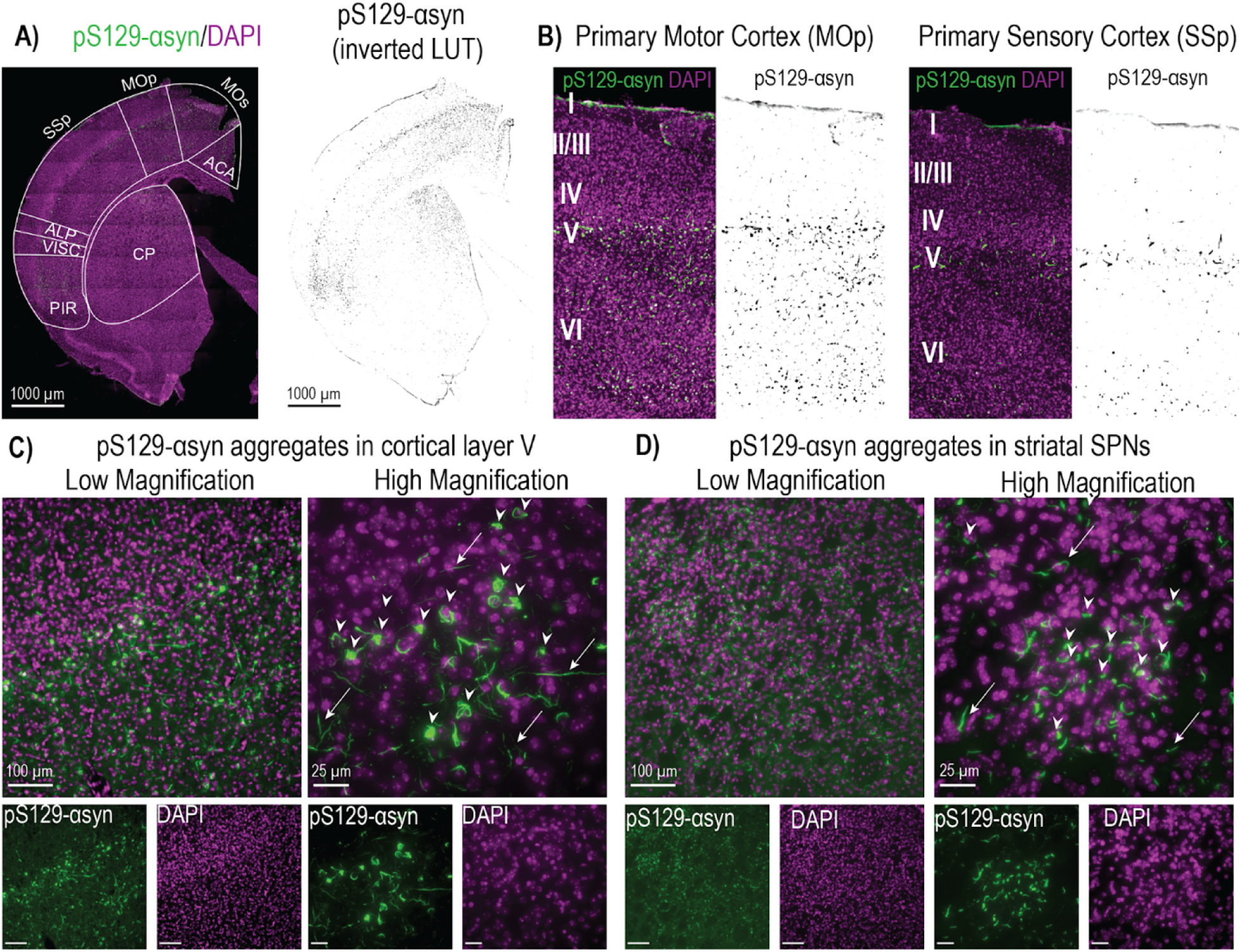
□-Syn aggregate formation in cortex and striatum at early time points post PFF inoculation. A) Immunofluorescence microscopy image of whole hemisphere of a 6-week striatally PFF-injected, representative animal stained for anti-pS129-□-syn (green) and DAPI (magenta). pS129-α-syn-positive aggregates are visible in secondary motor cortex (MOs), primary motor cortex (MOp), primary sensory cortex (SSp), Agranular insular area (ALP), Visceral Area (VISC), the piriform cortex (PIR) and the dorsal striatum (CP). Inverted, black and white p129 signal is shown for aggregate visualization throughout the hemisphere. Scale bar 1000 µm. The selected hemisphere represents a coronal section 0.2 mm AP from bregma according to Allen mouse brain atlas B) 20X magnification into MOp and SSp shows large and robust somal and neuritic aggregate accumulation in cortical layer V. Layer VI also showed abundant somal and neuritic pathology, while pathology in layer II/III and layer I is primarily neuritic. C) Low magnification and high magnification images of pS129-α-syn-positive aggregates in motor cortex layer V highlight accumulation of somal pS129-α-syn (arrow heads) and neuritic (arrows) pathology in this specific cortical layer. D) Low magnification and high magnification of pS129-α-syn-positive aggregates in dorsal striatum, arrowheads highlight somal pathology in spiny projection neurons and arrows point toward neuritic □-syn pathology.

### Excitatory synaptic density and morphology in the striatum six weeks following PFF injections

Using a combination of ExM and an Imaris 3D surface rendering (Fig S3a), corticostriatal synaptic loci in the dorsal striatum of either 6-weeks bilaterally PFF injected, or control (PBS and monomer) animals were imaged and quantified. To visualize corticostriatal synapses, the cortical, excitatory terminal marker VLGUT1, and the postsynaptic, excitatory marker Homer1 were used (Fig 2a). Total surfaces were reconstructed from confocal z-stacks using Imaris imaging software and synaptic loci were isolated by filtering synaptic surfaces in close proximity to the juxtaposed synaptic partner (<0.01µm). No change between groups was observed for the density of total Homer1 surfaces in the dorsal striatum, but density of total VGLUT1 surfaces was significantly reduced in PFF-injected animals compared to controls (Fig 2b). The density of corticostriatal synaptic loci for VLGUT1 and Homer1 were both significantly reduced in mice six weeks following injections with PFFs (Fig 2b). PFF-injected animals also showed significant decreases in volumes of total and synaptic VGUT1 and total Homer1, compared to controls (Fig 2c), while no change in volume overlap of synaptic loci with their juxtaposed synaptic partner was observed between groups (Fig 2c). Approximately 80% of total VGLUT1 (78.69±3.81% for control group) surfaces were characterized as corticostriatal synaptic loci, and the percentage of synaptic loci of total VGLUT1 surface count was not different between groups (Fig 2d). Around 20% of Homer1 (22.98±02.59% for control group) surfaces were classified as corticostriatal synaptic loci for control animals in the dorsal striatum, however this percentage was significantly reduced in PFF-injected animals (Fig 2d).

**Figure 2:**
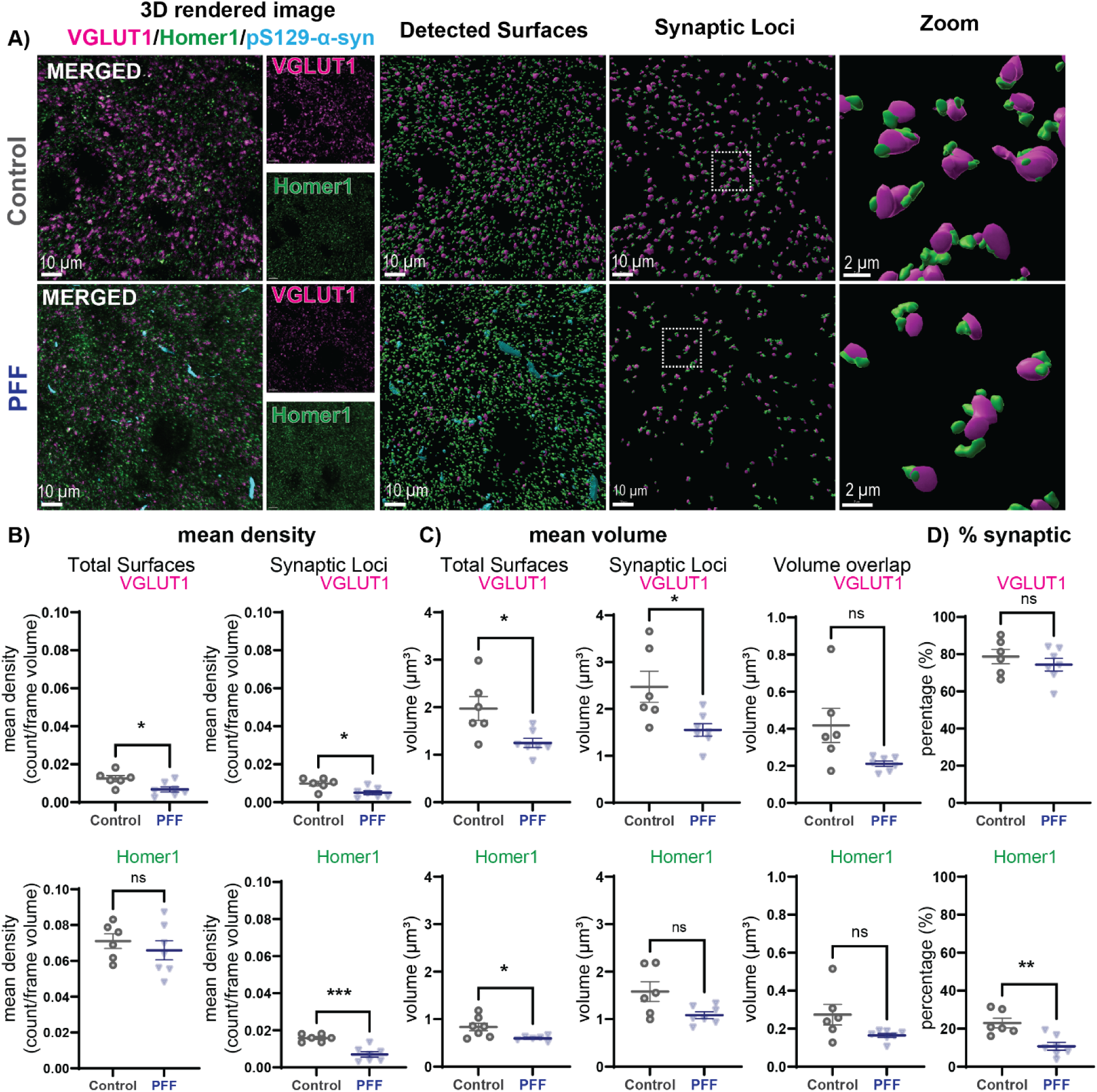
Striatal PFF injections causes reduction in VGLUT1/Homer1 synaptic loci in the striatum. A) From left to right: Left panel showing 3D-rendered confocal images of VGLUT1 (magenta), Homer1 (green) and pS129-α-syn (cyan) for control and PFF-injected animals. The next panel shows detected surfaces for the markers of interest. Panels on the right show filtered synaptic loci consisting of VGLUT1/Homer1 surfaces in close proximity with inset to show juxtaposed positioning of corticostriatal synaptic markers. B) Quantification of density of total VGLUT1 and Homer1 surfaces: Total density of VGLUT1 surfaces was significantly reduced (Unpaired Student‘s t-test, p=0.0194) in PFF-injected animals, while total Homer1 surface density did not change (Unpaired Student‘s t-test, p=0.467). Density of synaptic loci for VGLUT1 and Homer1 were both significantly decreased in pathology-harboring animals (Synaptic VGLUT1 density: unpaired Student‘s t-test, p=0.0108, synaptic Homer1 density: unpaired Student‘s t-test, p=0.0004). C) Quantification of corticostriatal synaptic surface volumes: total VGLUT1 surface volumes (Unpaired Student‘s t-test, p=0.0157) as well as synaptic VGLUT1 volumes (Unpaired Student‘s t-test, p=0.0196) were significantly decreased in PFF-injected animals, as well as total postsynaptic Homer1 volumes (Welch t-test, p=0.026). Volume overlap with juxtaposed surfaces did not differ between control and PFF animals. D) Percentage of synaptic loci for total surfaces was significantly reduced for postsynaptic Homer1 (Unpaired Student’s t-test, p=0.0032) in PFF animals, while % of synaptic VGLUT1 did not change (Unpaired Student’s t-test, p=0.4106). Statistical significance defined as: *p<0.05, **p<0.01, ***p<0.001, ****p<0.0001***. Corresponding statistical analysis information and group mean□SEM are provided in table 3.

**Table 3:**
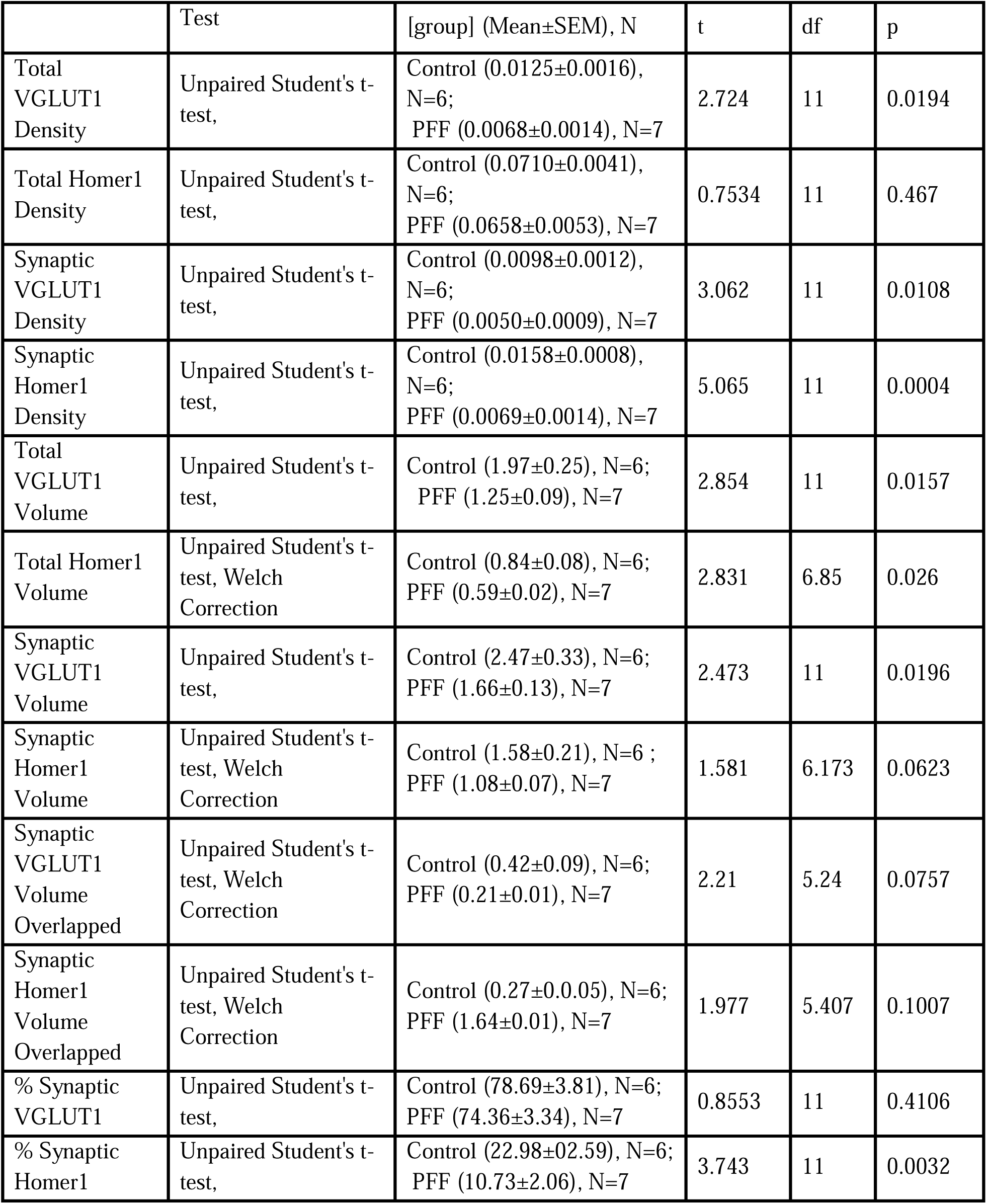
Statistics Summary Table for Figure 2.

**Table 4:** Statistics Summary Table for Figure 3. *: Outlier removed (Grout test)

|  | Test | [group] (Mean±SEM), N | t | df | p |
| --- | --- | --- | --- | --- | --- |
| %w pS129 in Homer1 vs VGLUT1 | Unpaired Student's t-test, Welch Correction | Homer1 (2.72±0.52), N=6*; VGLUT1 (18.92±3.58), N=7 | 4.483 | 6.253 | 0.038 |
| Synaptic VGLUT1 Volume | Unpaired Student's t-test, Welch Correction | wo pS129 (1.18±0.11), N=7; w pS129 (3.01±0.37), N=6 | 4.788 | 7.042 | 0.002 |
| Synaptic VGLUT1 Volume Overlapped | Unpaired Student's t-test | wo pS129 (0.17±0.02), N=7; w pS129 (0.42±0.03), N=7 | 7.159 | 12 | <0.0001 |
| Synaptic Homer1 Volume | Unpaired Student's t-test | wo pS129 (1.06±0.08), N=7; w pS129 (1.91±0.16), N=6* | 5.013 | 11 | 0.0004 |
| Synaptic Homer1 Volume Overlapped | Mann-Whitney test | wo pS129 (0.14±0.01), N=7; w pS129 (0.60±0.09), N=7 |  |  | 0.0006 |

To validate the findings from ExM experiments, the same analysis pipeline was applied to brain samples of the same animal cohort processed using confocal microscopy. Similar to ExM results, pS129-α-syn pathology is only observed in PFF-injected animals, while the negative control groups (PBS and Monomer injections) show no inclusions (Fig S4a). ANOVA statistical analysis revealed significantly reduced VGLUT1 density in PFF-injected animals compared to PBS and Monomer control groups (Fig S4b), while no statistical differences between the negative control groups were observed. ANOVA analysis of total Homer1 density revealed no statistical difference between PFF-injected or control-injected groups. These findings match the results obtained by the ExM protocol, confirming the early loss of corticostriatal synapses upon pathology formation using an alternative protocol.

To assess whether pathological α-syn affects other excitatory inputs in the striatum, the same parameters as described in Figure 2 were assessed for thalamostriatal synapses. Thalamostriatal synapses were identified by staining for the excitatory, thalamic terminal marker VGLUT2 and excitatory, postsynaptic marker Homer1 (Fig S5). VGLUT2 fluorescent signal was lost during ExM protocol, thus samples were processed using standard immunofluorescence and confocal imaging. No change in density in thalamostriatal synaptic loci nor changes in total surfaces for Homer1 and VGLUT2 were observed between groups at 6-weeks post injections (Fig S5b, c). There was a significant increase in total and synaptic VGLUT2 surface volumes in PFF-injected animals compared to PBS-control animals (Fig S5c). Thus, reduction of excitatory synaptic loci caused by early α-syn aggregation occurs for corticostriatal but not thalamostriatal synapses. without change in thalamostriatal synaptic loci suggests a selective, pathological impact of toxic α-syn species on corticostriatal circuitry.

### Changes to synaptic morphology in synapses with p-α-syn aggregates

Using an object based colocalization approach in Imaris software, synaptic loci for VGLUT1 and Homer1 were separated into surfaces with and without pS129-α-syn in the dorsal striatum of mice six-weeks post bilateral PFF (Fig 3a). Almost 20% of presynaptic VGLUT1 synaptic loci (18.92±3.58%) colocalized with α-syn pathology, while only a small proportion of postsynaptic, Homer1 positive, corticostriatal loci markers (2.72±0.52%) showed colocalization with p-S129-α-syn aggregates (Fig 3b). Both synaptic VGLUT1 and Homer1 with p-S129-α-syn aggregates showed a significant increase in volume compared to those without p-S129-α-syn (Fig 3c). A significant increase in volume overlap with juxtaposed synaptic loci partners for VGLUT1 with p-S129-α-syn and Homer1 with p-S129-α-syn was also observed. To assess whether synaptically-localizing p-S129-α-syn had similar effects on thalamostriatal synapses, a similar approach of object-based colocalization of p-S129-α-syn aggregates in VGLUT2/Homer1 synaptic loci was performed in the dorsal striatum (Fig S6). While only a small percentage of VGLUT2 (5.24±0.77%) synaptic loci overlapped with pS129-α-syn, a significant increase in volumes in synaptic VGLUT2 surfaces positive for pS129-α-syn (VGLUT2 w pS129, Fig S6c) was observed. Furthermore, a significant increase in volume of VGLUT2 juxtaposed Homer1 synaptic loci positive for pS129-α-syn was observed in the dorsal striatum (Fig S6c). While a larger percentage of corticostriatal synapses presented positive with pS129-α-syn pathology (18.92±3.58%) as opposed to thalamostriatal synapses (5.24±0.77%), these findings point to a critical impact of small aggregates on excitatory synapse morphology in the striatum at an early timepoint post pathology initiation.

**Figure 3:**
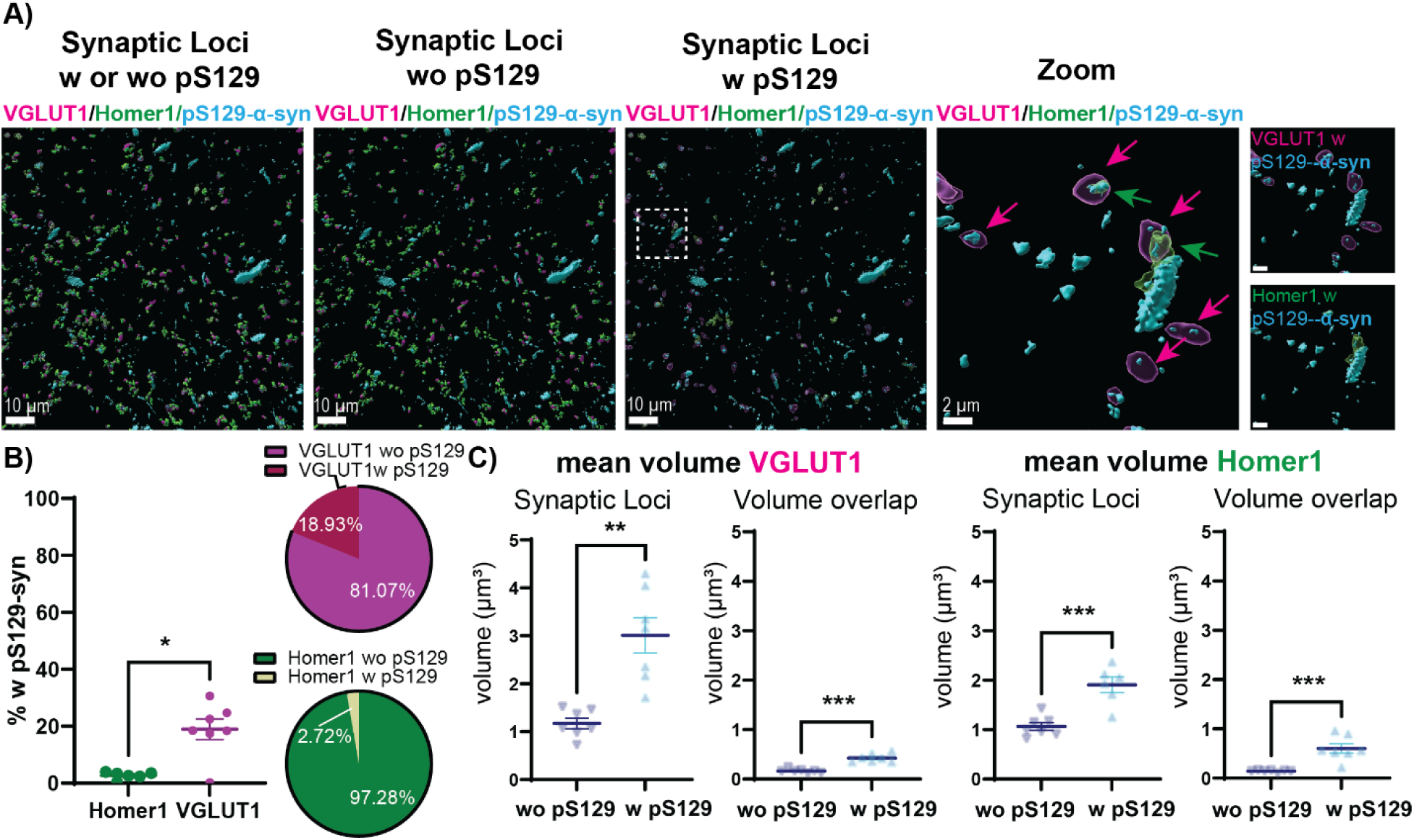
Morphological changes to pS129-α-syn positive synaptic compartments. A) Shown are reconstructed, filtered corticostriatal synaptic loci (VGLUT1-Homer1) and pS129-α-syn pathology in the dorsal striatum of animals 6-weeks post PFF injections. Subsets of VGLUT1 and Homer1 synaptic loci showed co-localization with pS129-α-syn aggregates and are highlighted as transparent surfaces. Additional filtering allowed for the display of synaptic loci without (wo pS129) pS129-α-syn aggregates and synaptic loci with (w pS129) pS129-α-syn aggregates. A zoom-in of synaptic loci w pS129 highlights synaptic VGLUT1 w pS129 (magenta arrow and surface) and synaptic Homer1 w pS129 (green arrow and surface), pathologic pS129-α-syn aggregates are shown in cyan. B) Quantification of percentage of synaptic loci with pS129-α-syn aggregates: VGLUT1 synaptic loci showed a significantly higher positive percentage for small, intrasynaptic pS129-α-syn aggregates than Homer1 synaptic loci. (Welch t-test, p=0.038). Pie charts for visualization of % w/wo pS129 for the respective corticostriatal synaptic loci VGLUT1 (magenta) and Homer1 (green) are shown on the right. C) Volumes of synaptic loci positive for pS129-aggregates (w pS129) show a significant enlargement in volumes for VGLUT1 (Welch t-test, p=0.002) and Homer1 (Unpaired Student’s t-test, p<0.0001). Volume overlapped with respective synaptic partners of synaptic loci w pS129 were significantly larger compared to synaptic loci wo pS129 (VGLUT1 volume overlapped: unpaired Student’s t-test, p=0.0004, Homer1 volume overlapped: Mann-Whitney test, p=0.0006). Statistical significance defined as: *p<0.05, **p<0.01, ***p<0.001, ****p<0.0001***. Corresponding statistical analysis information and group mean□SEM are provided in table 3.

### Expression of synaptic proteins in the striatum six weeks following PFF-induced α-syn aggregate formation

Expression of synaptic proteins was quantified in the striatum from control (PBS) mice and mice six weeks following PFF injections (Fig 4). Levels of VGLUT1 were significantly reduced in the striatum of PFF-injected mice compared to controls. Conversely, expression of VGLUT2 was significantly increased in PFF-injected animals. Levels of TH expressed in dopaminergic fibers in the striatum were not reduced in PFF-injected mice compared to controls. Protein expression levels of SNARE proteins, Snap25 and VAMP2 in the dorsal striatum of pathology-harboring mice were significantly reduced in this present study. There were no significant reductions in post-synaptic Homer1 in the striatum of PFF-injected mice compared to controls. No changes in protein expression levels of the AMPAR subunits GluA1 and GluA2 were observed, but protein expression of GluA3, a subunit required for basal synaptic transmission, was significantly reduced in the striatum of PFF-injected animals compared to controls. There was also a significant reduction in NMDAR2b subunit, while NMDAR1 and NMDAR2a levels were not significantly reduced in PFF-injected animals. Overall, these data show selective reduction of presynaptic cortico-striatal proteins with specific reductions in post-synaptic ionotropic receptor density. Immunoblot protein quantification revealed a significant impact of pathological α-syn on synaptic protein expression in the striatum, pointing to a critical connection of synaptic integrity and α-syn pathology.

**Figure 4:**
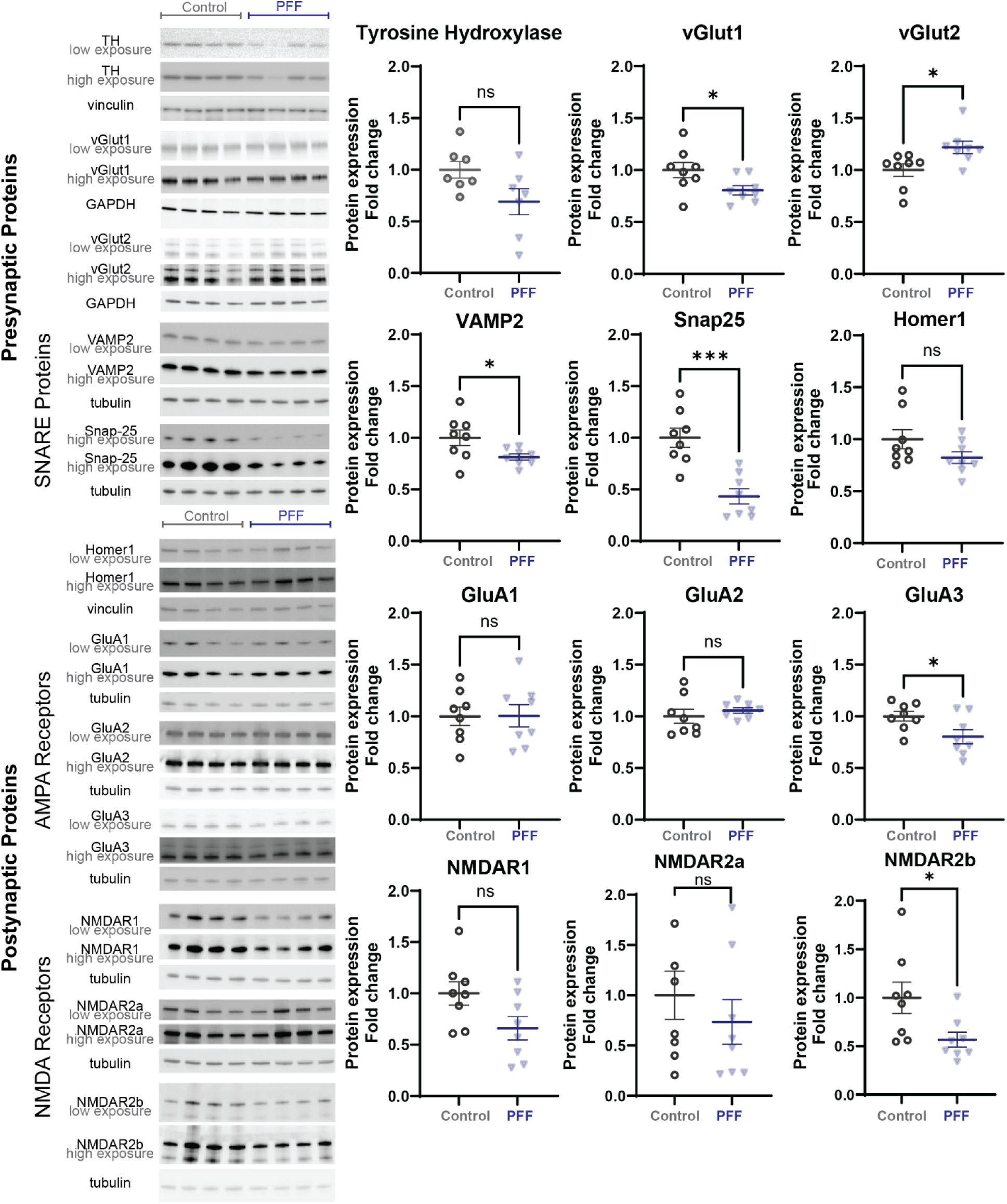
Changes in synaptic protein expression in the striatum of PFF-injected animals. Immunoblotting of Triton-X soluble striatal protein lysates of PFF-or PBS-treated (control) animals 6-weeks post-injections. Low and high exposure of chemiluminescent or fluorescent protein blots for markers of interest are shown as well as their respective loading control (vinculin, tubulin, or GAPDH). Protein expression is quantified as fold change compared to the control group. Presynaptic markers VGLUT1 (Unpaired Student’s t-test, p=0.0386) and SNARE proteins VAMP2 (Unpaired Student’s t-test, p=0.047) and Snap25 (Unpaired Student’s t-test, p=0.0003) were significantly reduced in the striatum of PFF-injected animals, while VGLUT2 (Mann-Whitney test, p=0.0104) expression was increased. A significant reduction in postsynaptic protein expression levels were observed for the AMPAR subunit GluA3 (Unpaired Student’s t-test, p=0.031) and NMDAR subunit NMDAR2b (Unpaired Student’s t-test, p=0.0291) in PFF injected animals. Statistical significance defined as: *p<0.05, **p<0.01, ***p<0.001, ****p<0.0001***. Corresponding statistical analysis information and group mean□SEM are provided in table 5.

**Table 5:** Statistics Summary Table for Figure 4.

|  | Test | [group] (Mean±SEM), N | t | df | p |
| --- | --- | --- | --- | --- | --- |
| TH | Unpaired Student's t-test | PBS Control (1.00±0.08), N=7; PFF (0.69±0.13), N=7 | 2.07 | 12 | 0.0607 |
| VGLUT1 | Unpaired Student's t-test | PBS Control (1.00±0.06), N=8; PFF (1.22±0.06), N=8 | 2.283 | 14 | 0.0386 |
| VGLUT2 | Mann-Whitney test | PBS Control (1.00±0.07), N=8; PFF (0.81±0.04), N=8 |  |  | 0.0104 |
| Vamp2 | Unpaired Student's t-test, Welch Correction | PBS Control (1.00±0.08), N=8; PFF (0.81±0.03), N=8 | 2.303 | 8.936 | 0.047 |
| Snap25 | Unpaired Student's t-test | PBS Control (1.00±0.09), N=8; PFF (0.43±0.07), N=8 | 4.805 | 14 | 0.0003 |
| Homer1 | Unpaired Student's t-test | PBS Control (1.00±0.09), N=8; PFF (0.82±0.06), N=8 | 1.625 | 14 | 0.1264 |
| GluA1 | Unpaired Student's t-test | PBS Control (1.00±0.09), N=8; PFF (1.01±0.11), N=8 | 0.03451 | 14 | 0.973 |
| GluA2 | Unpaired Student's t-test, Welch Correction | PBS Control (1.00±0.07), N=8; PFF (1.06±0.03), N=8 | 0.7747 | 8.97 | 0.4584 |
| GluA3 | Unpaired Student's | PBS Control (1.00±0.05), | 2.399 | 14 | 0.031 |
|  | t-test | N=8; PFF (0.80±0.05), N=8 |  |  |  |
| NMDAR1 | Unpaired Student's t-test | PBS Control (1.00±0.11), N=8; PFF (0.65±0.11), N=8 | 2.133 | 14 | 0.0511 |
| NMDAR2a | Mann-Whitney test | PBS Control (1.00±0.23), N=8; PFF (0.73±0.22), N=8 |  |  | 0.5054 |
| NMDAR2b | Unpaired Student's t-test | PBS Control (1.00±0.16), N=8; PFF (0.56±0.08), N=8 | 2.43 | 14 | 0.0291 |

### Corticostriatal glutamate drive decreases in the presence of pathologic α-syn

Templated formation of α-syn aggregates has been shown to cause dysfunction of excitatory synapses. Due to the robust abundance of pathology in striatally-projecting cortical layer V neurons (Fig 1), the observed significant reduction in corticostriatal synaptic density (Fig 2) and morphological abnormalities in PFF animals (Fig 3), we examined how corticostriatal synaptic physiology is affected by the presence of early pathological α-syn. Whole cell patch clamp recordings of dorsal striatum spiny projection neurons were performed to assess total corticostriatal glutamate drive in 6-weeks post bilateral, striatal PFF-injected animals, as well as a monomer-control group. Using electrical stimulation of the corpus callosum to elicit neurotransmitter release from cortical terminals onto SPNs, evoked excitatory postsynaptic currents (EPSC) in response to increasing CC stimulation was measured. A mixed effect analysis revealed that PFF-treated animals showed significantly decreased evoked cortical glutamate transmission onto SPNs compared to control (monomer) injected animals at higher CC stimulation intensities (Fig 5b). Quantification of the paired pulse ratio from paired-pulse CC stimulation experiments showed no changes between PFF and monomer injected animals (Fig 5c). Recording and analyzing spontaneous EPSCs to assess overall excitatory synaptic transmission onto SPNs did not show significant differences between PFF and monomer injected animals. Post Hoc staining of coronal sections from physiological recordings revealed robust cortical pathology, accumulating in layer V and IV (Fig S7), further suggesting that pathological α-syn might contribute to the observed synaptic dysfunction in basal transmission. These results suggest critical physiological dysfunction of corticostriatal synapses in the presence of pathological α-syn.

**Figure 5:**
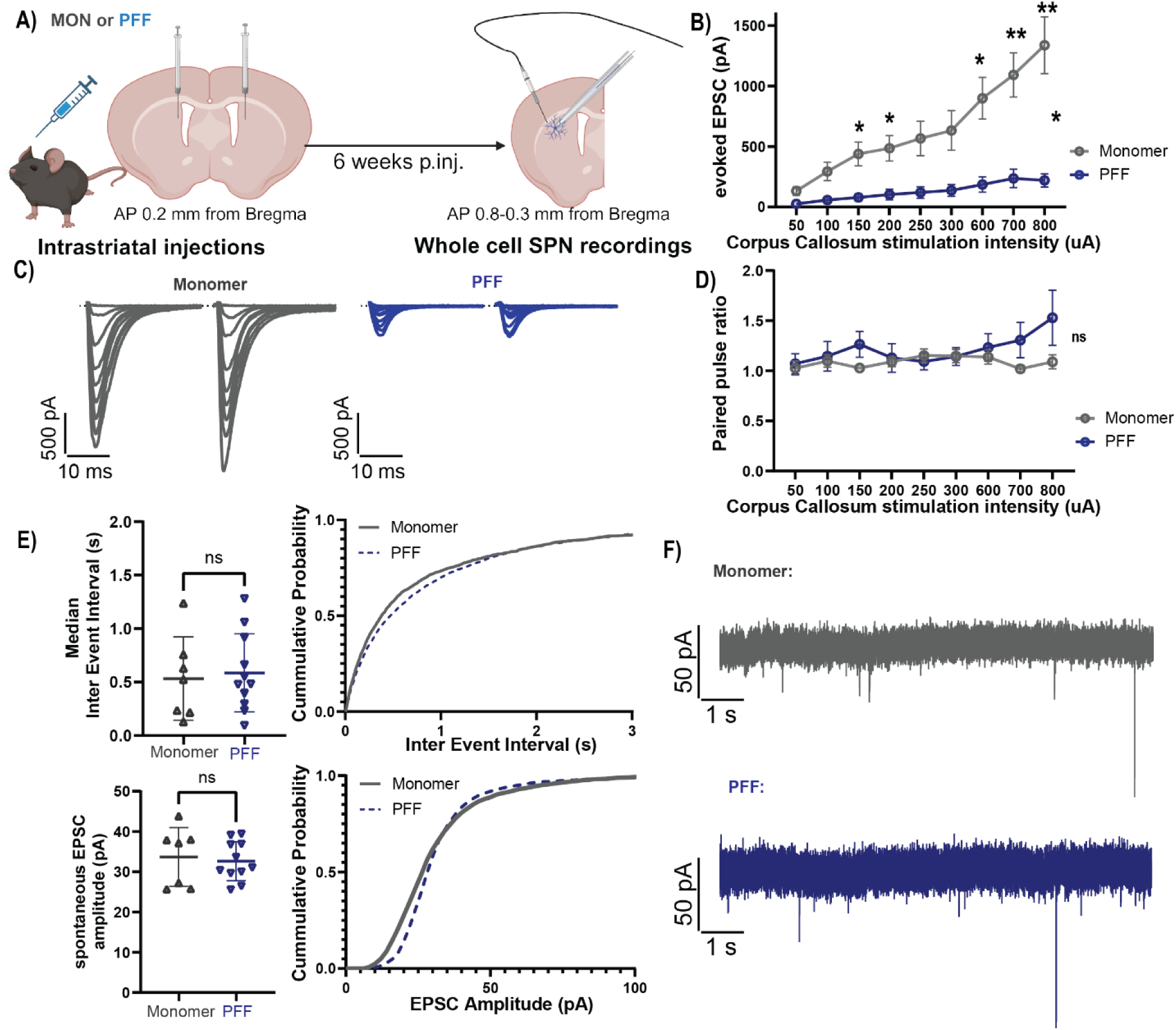
Decreased corticostriatal glutamate drive in mice harboring. □**-syn pathology.** A) Schematic illustration of experimental setup: 3-month-old C57BL/6 mice received bilateral injections of PFFs or monomeric □-syn as control and whole-cell patch clamp recordings of SPNs in close proximity to the corpus callosum (CC) were performed 6-weeks post-injections. B) Input-output curve to assess total corticostriatal glutamate drive by stimulating the CC with increasing intensity and measuring the evoked postsynaptic current in SPNs. PFF-injected animals showed a significant reduction in total cortex-to-striatum, evoked EPSC compared to control (Mixed-effect analysis with Geisser-Greenhouse correction, Šidák’s multiple comparison post hoc). C) Quantification of paired pulse ratio measured by paired pulse stimulation (75 ms interval) of the CC and evoked EPSC showed no significant difference between groups (Mixed-effect analysis with Geisser-Greenhouse correction). D) Representative EPSC traces for CC paired pulse stimulation with increasing stimulus intensity for control-treated animals (gray) and PFF treated animals (blue). E) Quantification of total spontaneous synaptic activity on SPNs in control and PFF-treated animals. Quantification of median inter event interval (IEI) and mean amplitude of sEPSCs and cumulative probabilities are shown. No differences were observed between groups (IEI: Unpaired Student‘s t-test: p=0.771, amplitude: Unpaired Student‘s t-test: p=0.7208). Cumulative probability analysis of the inter event interval points to a decrease in overall synaptic activity from recordings in PFF-injected animals. F) Representative 10 second traces of voltage clamp recordings of SPNs showing spontaneous EPSCs for both treatment groups are shown. Statistical significance defined as: *p<0.05, **p<0.01, ***p<0.001, ****p<0.0001***. Corresponding statistical analysis information and group mean□SEM are provided in table 6.

**Table 6:** Statistics Summary Table for Figure 5.

|  | Test | [group] (Mean±SEM), n, N | F | Dfn | Dfd | p |
| --- | --- | --- | --- | --- | --- | --- |
| Evoked EPSC Repeated Measure | Mixed-effect analysis with Geisser-Greenhouse correction | Monomer, n=13, N=7;<br>PFF, n=14, N=8;<br>Row Factor;<br>Column Factor;<br>Row Factor x Column Factor | 20.49<br>20.43<br>9.049 | 2.103<br>1<br>8 | 52.31<br>25<br>199 | <0.0001<br>0.0001<br><0.0001 |
| Evoked EPSC 50 μA | Šidák's multiple comparison post hoc | Monomer (132.6±35.3), n=13, N=7;<br>PFF (26.3±8.1), n=14, N=8 | | | | 0.0985 |
| Evoked EPSC 100 μA | Šidák's multiple comparison post hoc | Monomer (294.5±76.9), n=13, N=7;<br>PFF (57.7±23.2), n=14, N=8 | | | | 0.0905 |
| Evoked EPSC 150 μA | Šidák's multiple comparison post hoc | Monomer (438.9±97.6), n=13, N=7;<br>PFF (78.7±29.7), n=14, N=8 | | | | 0.029 |
| Evoked EPSC 200 μA | Šidák's multiple comparison post hoc | Monomer (486.3±105.2), n=13, N=7;<br>PFF (102.9±44.1), n=14, N=8 | | | | 0.0349 |
| Evoked EPSC 250 μA | Šidák's multiple comparison post hoc | Monomer (567.5±142.6), n=13, N=7;<br>PFF (120.1±47.2), n=14, N=8 | | | | 0.0803 |
| Evoked EPSC 300 μA | Šidák's multiple comparison post hoc | Monomer (632.8±163.6), n=13, N=7; PFF (137.5±48.2), n=14, N=8 | | | | 0.0987 |
| Evoked EPSC 600 μA | Šidák's multiple comparison post hoc | Monomer (899.6±172.8), n=13, N=7; PFF (185.9±63.1), n=14, N=8 | | | | 0.0131 |
| Evoked EPSC 700 μA | Šidák's multiple comparison post hoc | Monomer (1092.6±183.3), n=13, N=7; PFF (263.8±75.5), n=14, N=8 | | | | 0.0048 |
| Evoked | Šidák's multiple | Monomer (13338.2±234.7), |  |  |  | 0.0039 |
| EPSC 800 μA | comparison post hoc | n=13, N=7; PFF (220.1±54.4), n=13, N=8 | | | | |
| Paired-Pulse Ratio Repeated Measure | 2-Way RM ANOVA with Geisser-Greenhouse correction | Monomer, n=13, N=7; PFF, n=14, N=8; Row Factor Column Factor Row Factor x Column Factor | 1.138<br>0.8820<br>1.966 | 8<br>3.290<br>1 | 200<br>82.26<br>25 | 0.3396<br>0.4619<br>0.1732 |
| | Test | [group] (Mean±SEM), n, N | | t | df | p |
| sEPSC Inter Event Interval | Unpaired Student's t-test | MEDIAN Monomer (0.53±0.15), n=7, N=4; PFF (0.59±0.11), n=10, N=5 | | 0.295<br>1 | 16 | 0.7717 |
| sEPSC Amplitude | Unpaired Student's t-test | Monomer (-33.7±2.8), n=7, N=4; PFF (-32.6±1.5), n=10, N=5 | | 0.363<br>7 | 16 | 0.7208 |

### SPNs show aberrant intrinsic excitability in animals harboring α-syn pathology

SPN activity and excitability is modulated by dopamine, and dopaminergic denervation studies have been shown to affect the intrinsic properties of SPNs. However, how striatal α-syn aggregates contribute to SPN function and properties has not received attention. To assess whether the presence of pathological α-syn affects intrinsic SPN function, whole cell patch clamp recordings of SPNs were performed in the dorsal striatum of animals 6-weeks post bilateral, intrastriatal PFF and monomer control injections. Using the rheobase current as a measure of intrinsic excitability, a significant increase in rheobase current in SPNs in the PFF-injection cohort (Fig 6a) was observed. PFF injections also caused a significant increase in threshold membrane potential, with increased membrane depolarization required to elicit action potentials in recorded SPNs (Fig 6b). 2-way RM ANOVA analysis revealed a significant reduction in evoked AP count upon depolarization in SPNs in the PFF group. The relationship between action potential count and depolarizing current injections showed a right shift in PPF-injected animals, further supporting the findings of decreased intrinsic excitability of SPNs in the presence of α-syn pathology in the mouse brain (Fig 6c). The differences in spike threshold and rheobase current between PFF and control cohort were not reflective of changes in intrinsic membrane properties as no divergence in resting membrane potential nor membrane resistance was observed between groups (Fig S8).

**Figure 6:**
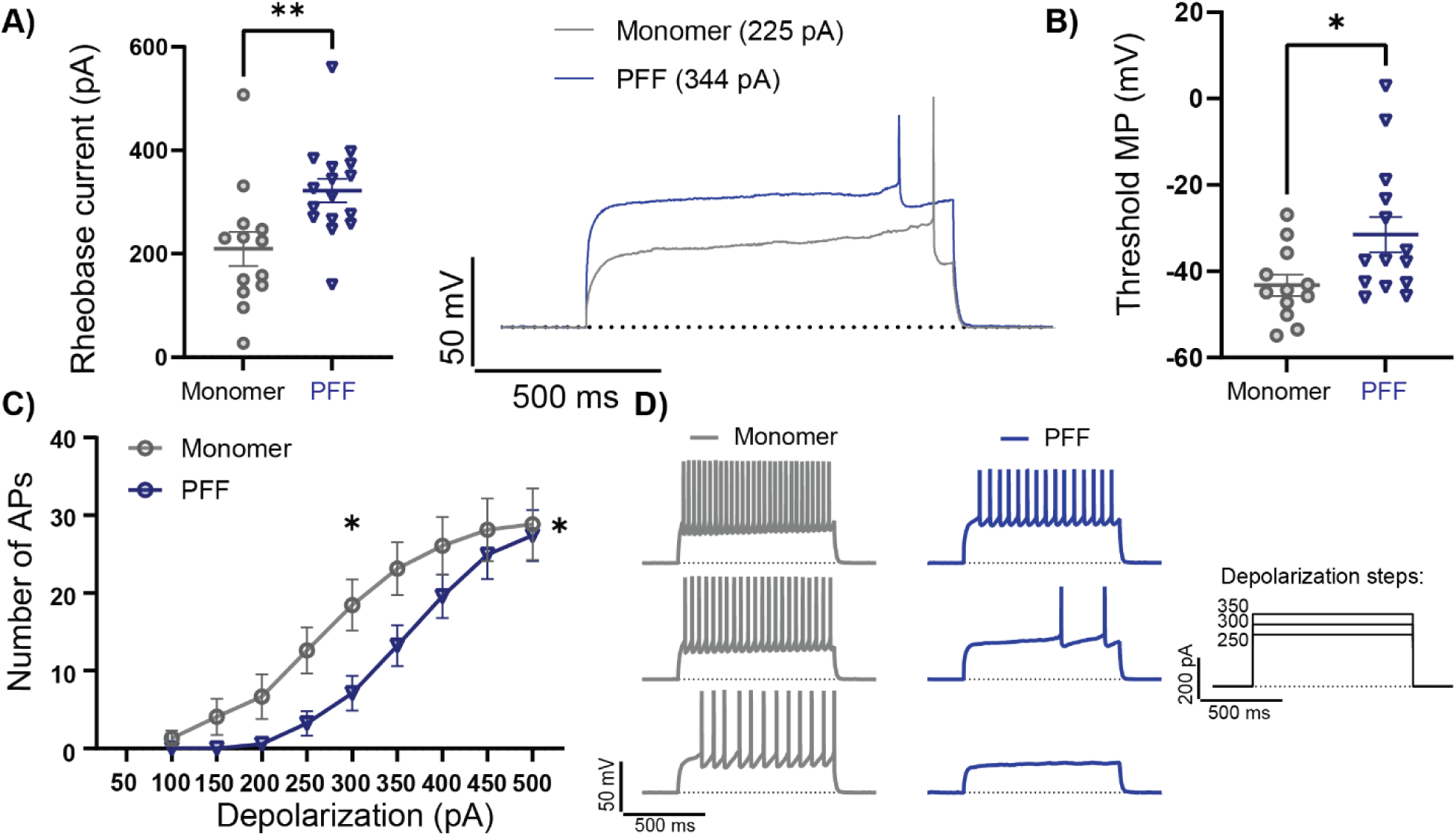
Decreased intrinsic excitability of SPNs in PFF treated animals. Whole cell patch clamp recordings were performed in the dorsal striatum of animals 6-weeks after receiving striatal PFF or control monomer injections A) SPNs of PFF treated animals showed a significant increase in their respective rheobase current compared to control (Unpaired Student‘s t-test: p=0.0077). Example traces for PFF (blue, 344 pA) and monomer control (gray, 225 pA) rheobase action potentials are shown in the dle. B) Threshold membrane potential (MP) for firing APs was significantly higher in PFF-injected animals than monomer controls (Mann-Whitney test: p=0.0221). C) Graph of action potential counts upon 1 ms depolarizing current injections shows a significant main effect of PFF treatment on evoked AP counts (2-Way RM ANOVA, Column Factor: F(1, 25)=5.694, p=0.0001), while a reduction in action potential counts for the 300 pA depolarizing current injection steps in PFF injected animals compared to controls was observed (Šidák’s multiple comparison post hoc test: p=0.0279). D) Representative voltage traces and action potential firing of SPNs upon increasing current stimulation (250 pA, 300 pA and 350 pA, bottom to top) for monomer control (gray) and PFF injected animals (blue) are shown. Statistical significance defined as: *p<0.05, **p<0.01, ***p<0.001, ****p<0.0001***. Corresponding statistical analysis information and group mean□SEM are provided in table 7.

**Table 7:** Statistics Summary Table for Figure 6.

|  |  |  |  |  |  |  |
| --- | --- | --- | --- | --- | --- | --- |
| | Test | [group] (Mean±SEM), N | t | t | df | p |
| Rheobase current | Unpaired Student's t-test | Monomer (209.7±33.3), n=13, N=7; PFF (322.3±22.8), n=16, N=8 | | 2.877 | 27 | 0.0077 |
| Threshold Membrane potential | Mann-Whitney test | Monomer (-43.2±2.4), n=12, N=7; PFF (-31.9±3.8), n=15, N=8 | | | | 0.0221 |
| Test | | [group] (Mean±SEM), N | F | Dfn | Dfd | p |
| Number of APs | 2-Way RM ANOVA | Monomer, n=13, N=7; PFF, n=17, N=9<br>Row Factor<br>Column Factor<br>Row Factor x Column Factor | 49.10<br>5.694<br>1.468 | 8<br>1<br>8 | 52.31<br>25<br>199 | <0.0001<br>0.0001<br>0.1699 |
| Number of APs 100 pA | Šidák's multiple comparison post hoc | Monomer (1.31±0.97), n=13, N=7 PFF (0), n=17, N=9 | | | | >0.999 |
| Number of APs 150 pA | Šidák's multiple comparison post hoc | Monomer (4.08±2.31), n=13, N=7 PFF (0), n=17, N=9 | | | | 0.951 |
| Number of APs 200 pA | Šidák's multiple comparison post hoc | Monomer ($6.69 \pm 2.87$), $n=13$, $N=7$; PFF ($0.59 \pm 0.59$), $n=17$, $N=9$ | | | | 0.6489 |
| Number of APs 250 pA | Šidák's multiple comparison post hoc | Monomer ($12.62 \pm 2.98$), $n=13$, $N=7$; PFF ($3.24 \pm 1.58$), $n=17$, $N=9$ | | | | 0.1217 |
| Number of APs 300 pA | Šidák's multiple comparison post hoc | Monomer ($18.46 \pm 3.27$), $n=13$, $N=7$; PFF ($7.12 \pm 2.23$), $n=17$, $N=9$ | | | | 0.0279 |
| Number of APs 350 pA | Šidák's multiple comparison post hoc | Monomer ($23.15 \pm 3.41$), $n=13$, $N=7$; PFF ($13.24 \pm 2.62$), $n=17$, $N=9$ | | | | 0.0836 |
| Number of APs 400 pA | Šidák's multiple comparison post hoc | Monomer ($26.08 \pm 3.71$), $n=13$, $N=7$; PFF ($19.59 \pm 2.81$), $n=17$, $N=9$ | | | | 0.5688 |
| Number of APs 450 pA | Šidák's multiple comparison post hoc | Monomer ($28.15 \pm 4.03$), $n=13$, $N=7$; PFF ($24.94 \pm 3.13$), $n=17$, $N=9$ | | | | 0.9898 |
| Number of APs 500 pA | Šidák's multiple comparison post hoc | Monomer ($28.85 \pm 4.61$), $n=13$, $N=7$; PFF ($27.41 \pm 3.28$), $n=17$, $N=9$ | | | | 0.9898 |

### Early α-syn aggregation is accompanied by dopaminergic denervation

Dopaminergic denervation of the caudate and putamen (homologous to the striatum in mice) has been well characterized in PD and is thought to contribute to striatal dysfunction in motor behavior. Previous studies using striatal PFF injections in mice show dopaminergic (DA) denervation in the striatum and neural degeneration in the substantia nigra pars compacta (SNpc) as early as 3 months post injections. To assess whether the templated formation of α-syn pathology causes dopaminergic terminal loss at earlier time points, the immunofluorescence intensity of dopaminergic terminal marker tyrosine hydroxylase was quantified in 6-weeks post bilateral PFF-injected brains. Mean fluorescent intensity of immunofluorescent staining for neuronal marker NeuN was used to normalize TH fluorescent signal, as NeuN fluorescent intensity did not differ between treatment groups (Fig S9b). Early loss of tyrosine hydroxylase (TH) fibers in the dorsal striatum was observed in bilaterally injected C57BL/6 mice six weeks post pathology initiation compared to control-injected mice (Fig S9b). In conjunction with previously reported work, a significant reduction in dorsal striatum TH fiber density was observed 12 weeks post PFF injections compared to controls. However, there was no major effect of time point post injections on TH fiber density, with the 6-weeks post PFF injections already causing maximal dopaminergic denervation. Taken together, these results showed that early templated formation of α-syn pathology initiated in the mouse striatum significantly affects nigrostriatal circuits as early as 6-weeks post injections.

## Discussion

Striatal function is impaired in PD contributing to both the motor and non-motor symptoms of disease (Zhai et al., 2018). α-Syn inclusions are found in IT neurons of the cortex that project to the striatum, but how the presence of pathological α-syn aggregates on corticostriatal synaptic structure and function is relatively unexplored. Increasing evidence has linked α-syn pathology with excitatory synaptic dysfunction (Chen et al., 2022; Wu et al., 2019; Zhou et al., 2024), and here we rigorously elucidated the impact of α-syn aggregation on PD-relevant corticostriatal synaptic structure and function. In this present study, we show alterations in multiple electrophysiological, structural, and biochemical measures of synaptic function, which all point to loss of, and decreased function of corticostriatal projections harboring PFF-induced α-syn aggregates. Taken together, our findings suggest that corruption of synaptic α-syn into pathological forms induces synaptic dysfunction through multiple mechanisms quickly after becoming corrupted. These data suggest that some symptoms of PD may be due to synaptic dysfunction, particularly excitatory corticostriatal transmission, rather than overt neuron degeneration, which may be able to be targeted through increasing glutamatergic synaptic function beyond boosting dopamine levels.

Pathophysiological dysfunction of excitatory striatal synapses has been shown in dopaminergic lesion studies in rodents and monkeys, as well as post hum PD tissue. These studies have shown a role of dopamine loss in re-arranging multiple aspects of striatal physiology in PD (Graves and Surmeier, 2019). This dopamine centric view of PD however does not take into account the multiple cell types that develop LP, especially the large burden of pathology in cortical projection neurons (Goralski et al., 2024; Stoyka et al., 2020), and how pathological α-syn aggregates affect basic corticostriatal synaptic transmission. Our data reveals a significant reduction in total electrically evoked corticostriatal glutamate release onto SPNs, suggesting that α-syn reduces functionally connected cortico-striatal synapses or that α-syn aggregation decreases the efficacy of pre-synaptic corticostriatal synapses to release glutamate. Reduction in evoked glutamatergic transmission could be explained by numerous reasons. The presence of α-syn inclusions could change the overall release probability. However, we did not observe any changes in paired-pulse ratios between groups, presenting an argument against changes in pre-synaptic release probability. Furthermore, analysis of the frequencies of sEPSCs showed no changes in total excitatory, spontaneous synaptic transmission. However, sEPSCs recorded from SPNs represent total excitatory synaptic transmission, and it is not possible to distinguish between thalamic or cortical inputs. We therefore cannot rule out a compensatory increase in thalamostriatal glutamate release probability in response to a decrease in corticostriatal glutamate. Additionally, changes in synaptic strength or an overall decrease in the amount of active synaptic release sites could also contribute to the observed defects in evoked corticostriatal transmission. Our immunohistochemical findings showing decreased VGLUT1+ surfaces would suggest that this reduction in synapses may at least partially explain our decreased glutamate release findings. Additionally, we report decreased expression levels of striatal SNARE proteins essential for neurotransmitter release upon pathology formation, a finding supported by studies showing reduced SNARE protein expression in the presence of aggregated α-syn (Vallortigara et al., 2016; Volpicelli-Daley et al., 2011). Our findings are largely in agreement with other recent reports on the electrophysiological consequences of aggregation of α-syn. Recent reports have shown altered cortico-amygdala connectivity in the PFF model with decreased cortico-basolateral amygdala (BLA), but not thalamo-BLA, glutamate release (Chen et al., 2022; Zhou et al., 2024).

Downstream of changes in cortico-striatal glutamate release, we also find SPN intrinsic excitability to be significantly reduced in PFF injected animals. The presence of somal inclusions in SPNs could be a possible explanation of changes to intrinsic excitability due to altered protein homeostasis in aggregate-harboring neurons (Goralski et al., 2024). However, this would likely not influence all cells patched as the relative number of SPNs with somal inclusions is low compared to the total number of SPNs and may reflect compensatory changes in striatal activity due to altered total cortico-striatal drive. Finally, dopamine loss may partially drive our phenotype as we observe early dopaminergic terminal loss in the dorsal striatum in PFF-injected animals. This may additionally influence SPN excitability as DA denervation studies using 6-OHDA lesions have shown decreased excitability of D2 SPNs (Fieblinger et al., 2014). Corticostriatal synaptic transmission and SPN excitability are crucial parts of the basal ganglia motor circuitry, and synucleinopathy-related dysfunction through the mechanisms discussed above could contribute to behavioral symptoms seen in PD and DLB patients.

Our structural based imaging approach allowed us to characterize morphology of glutamatergic synapses that may be altered to cause the robust electrophysiological deficits that we observed. As expected, we saw a reduction in VGLUT1+ and Homer1+ corticostriatal synaptic loci and volume, suggesting degeneration of some of these synapses and altered function of remaining synapses. Overall reduction in synaptic protein expression of vGLUT1, Snap25 and VAMP2 in the striatum of PFF-injected mice support loss of synapses. However, postsynaptically, we only saw specific reductions in GluR3 and NMDAR2b suggesting a possible structural change in postsynaptic densities rather than an overall loss.

Interestingly, our data may suggest that this process is specific to corticostriatal synapses and not all glutamatergic synapses. We did not observe decreases in VGLUT2+ surfaces in our imaging approaches and observed elevated VGLUT2 in western blots of PFF injected tissue. This suggests that thalamic inputs into the striatum are not decreasing like cortical inputs, and that thalamic inputs may even be increasing in strength as a compensatory response to lost corticostriatal synapses. How this potential imbalance in cortical and thalamic glutamatergic inputs into the striatum alters basal ganglia function is not fully understood, but this imbalance may be an important driver of especially the motor symptoms of PD (Martel and Galvan, 2022). Our data would also suggest that this imbalance of cortical and thalamic inputs is an early consequence of aggregation of α-syn rather than an ongoing homeostatic change in response to DA loss or increasing α-syn aggregation. In context of the multiple animal model, post-mortem and *in vivo* imaging studies in DLB and PDD patients suggesting excitatory synapse dysfunction (Andersen et al., 2021; Froula et al., 2018; Sah et al., 2024; Stephens et al., 2005; Wilson et al., 2020; Wu et al., 2019), our data may point to a broader selective vulnerability of cortical synapses throughout the brain because in the basolateral amygdala, cortical, but not thalamic, excitatory synapses are impaired (Chen et al., 2022). These data may suggest that specifically boosting cortical, and not thalamic, function may be a novel way to symptomatically treat the motor and non-motor symptoms of α-synucleinopathies.

Paradoxical to the findings in reduced synaptic loci volumes in animals harboring pS129-α-syn pathology, we show increased volumes of reconstructed synaptic markers overlapping with small pS129-α-syn aggregates. The presence of synaptic α-syn aggregates could potentially cause a loss of organization or mislocalization of synaptic proteins which could result in the synapses appearing larger in volume. This could be a common synaptic mechanism as work from Gcwensa et al. on cortico-BLA and thalamo-BLA synapses, shows similar enlarged surface volumes when colocalized with pS129-α-syn (Gcwensa et al., 2024). Studies in human postmortem DLB brain also report enlargement of presynaptic compartments in close proximity to α-syn aggregates (Colom-Cadena et al.). In the amygdala, synaptic vesicles in PFF-injected mice are more clustered, which may cause increased overall synaptic volumes (Gcwensa et al., 2024). Our findings suggest that corruption of endogenous α-syn leads to rearrangement of the structure of corrupted synapses that eventually leads to them being susceptible to pruning or loss. Future studies will be needed to track if this increased volume and structural rearrangement lead to early synaptic loss after endogenous α-syn corruption.

Here, we provide evidence of early impairment of corticostriatal synaptic physiology and aberrant synaptic morphology in the presence of pathologic α-syn. Glutamatergic transmission in basal ganglia processing loops can affect can both movement and executive function associated information processing (Arrigoni et al., 2024; Dirnberger and Jahanshahi, 2013; Firbank et al., 2016; Fischer, 2021; Gómez-Ocádiz and Silberberg, 2023; Kemp et al., 2017), and α-syn aggregation directly disrupting these processing loops may represent a link between the manifestation of non-motor and motor symptoms present in α-synucleinopathies. Further understanding of how α-syn aggregation affects cortical transmission may lead to novel therapeutic strategies to treat symptoms of PD and DLB that are not currently addressed by current clinical gold standard of therapy for these disorders.

## List of Abbreviations

α-syn: α-synuclein
AMPAR: α-amino-3-hydroxy-5-methyl-4-isoxazoleproprionic acid Receptor
BLA: Basolateral Amygdala
DA: dopamine; dopaminergic
DLB: Dementia with Lewy Bodies
ExM: Expansion Microscopy
IB: Immunoblot
IT: Intratelenephalic
LBD: Lewy Body Disorders
LP: Lewy Pathology
MOp: Primary Motor Cortex
MOs: Secondary Motor Cortex
NMDAR: N-methyl-D-aspartate Receptor
PBS: Phosphate buffered Saline
PD: Parkinson’s Disease
PFF: Preformed Fibrils
PPR: Paired-pulse ratio
pS129-α-syn: phospho-serine 129 α-syn
Snap25: Synaptosomal-associated protein, 25 kDa
SNARE: Snap Receptor
SPN: Spiny Projection Neuron
SSp: Primary Somatosensory Cortex
TH: Tyrosine Hydroxylase
VAMP2: Vesicle associated membrane protein 2
VGLUT2: Vesicular Glutamate Transporter 1
VLGUT1: Vesicular Glutamate Transporter 2
w pS129: with Phospho-Serine 129 α-synuclein
wopS129: without Phospho-Serine 129 α-synuclein

## Author Contributions

CFB: Writing, Reviewing, Editing, Conceptualization, Investigation, Methodology, Formal Analysis, Data Curation, Visualization, Funding Acquisition

HC: Reviewing, Editing, Data Curation, Formal Analysis

NZG: Writing, Reviewing, Editing, Methodology

DH: Methodology, Data Curation, Reviewing

DN: Methodology, Data Curation, Reviewing

MM: Methodology, Data Curation, Reviewing

NC: Methodology, Data Curation, Reviewing

IG: Methodology, Data Curation, Formal Analysis, Reviewing

LVD: Supervision, Writing, Reviewing, Editing, Conceptualization, Investigation, Methodology, Funding Acquisition

MSM: Supervision, Writing, Reviewing, Editing, Conceptualization, Investigation, Methodology, Formal Analysis, Funding Acquisition

## Funding Information/Acknowledgements

The authors would like to acknowledge the following agencies and foundations for their generous support of our work: Parkinson Association of Alabama (predoctoral Fellowship to CFB), Parkinson’s Disease Foundation (Visiting Scholar Fellowship PF-VSF-1149805 and PF-VSF-944247 to CFB, MSM and LVD), Michael J. Fox Foundation (MJFF-023031 to MSM and LVD), NIH (R01NS132293, R01NS132293-S1, and R00NS110878 to MSM; R01AG081433 to LVD), the Harry T Mangurian Foundation (MSM), and the UF Fixel Institute Pilot Award (MSM).

## Declaration of generative AI in scientific writing

During the preparation of this work the authors used ChatGTP to help rephrase well-established material and methods sections (Animals, Preparation of α-syn preformed fibrils (PFF), Preparation of slices in Ex vivo Electrophysiology) After using ChatGTP, the authors reviewed and edited the content as need and take full responsibility of the content of the publication.

## Notes

### Competing Interest Statement

The authors have declared no competing interest.

