## Supplemental Figures for "Aggregated A-synuclein leads to corticostriatal synaptic dysfunction"

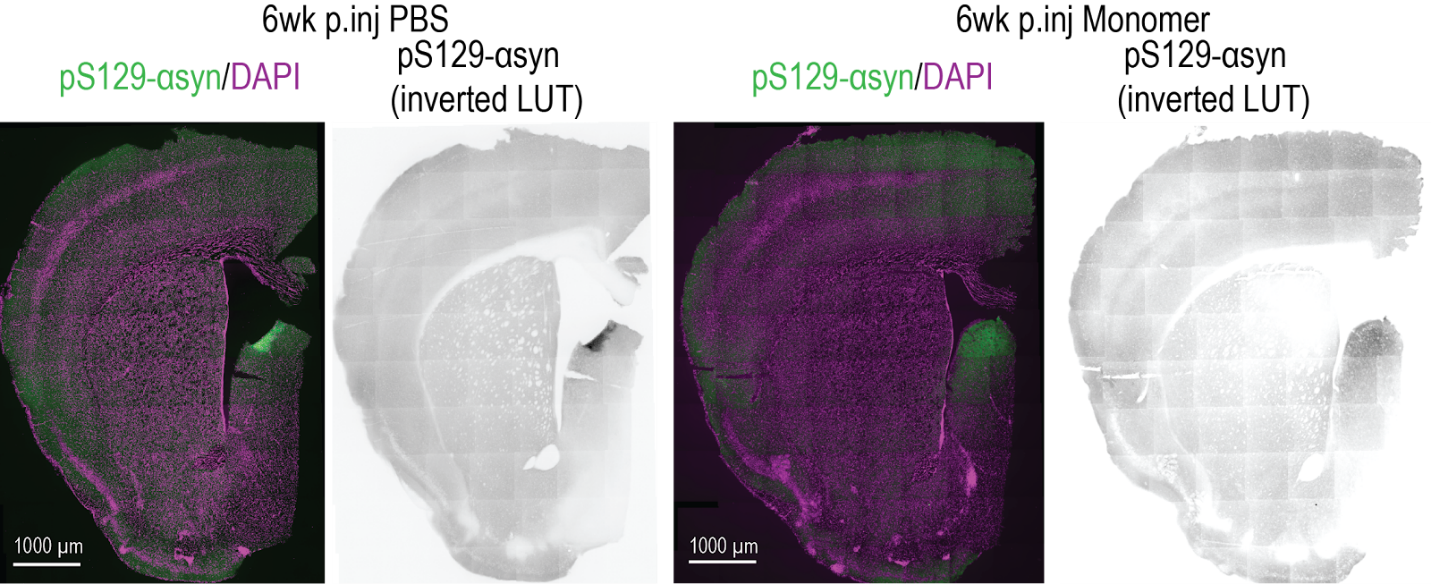

**Supplementary Figure 1: No presence of p129-positive aggregates in control injected animals.** Fluorescent widefield imaging of anti-p129-syn immunostainings (green) in control injected animals (PBS, monomeric α-syn) revealed an absence of pS129-α-syn-positive aggregates in striatum and cortex of control animals 6-weeks post injections.

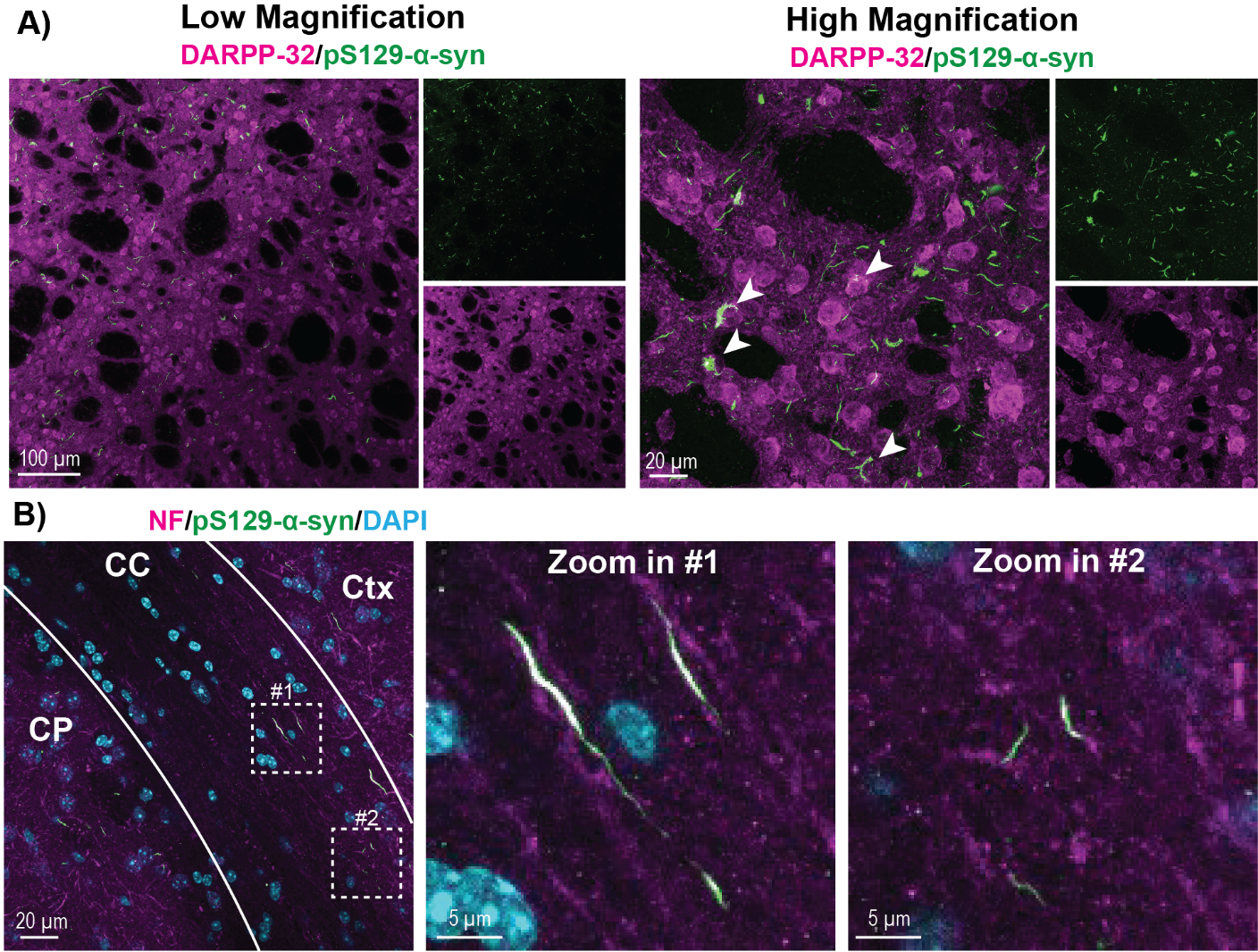

**Supplementary Figure 2: p129-positive aggregates can be found in collaterals of the corpus callosum and in SPNs in the striatum.** A) Low and high magnification fluorescent confocal imaging of anti-pS129-α-syn immunostainings (green) and SPN marker DARPP-32 (magenta) 6-weeks post striatal PFF injections revealed presence of somal pS129-α-syn-positive aggregates in SPNs, highlighted by white arrow heads. B) Confocal imaging of the CC 6-weeks post PFF injection reveals neuritic α-syn aggregates (green) colocalizing with axonal marker NF (magenta).

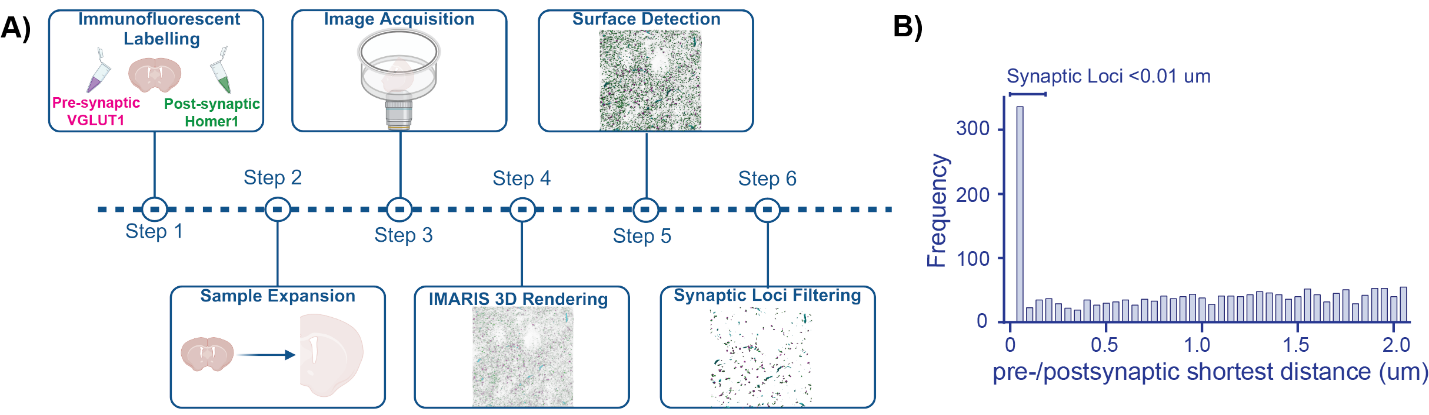
**Supplementary Figure 3: ExM and Imaris workflow to analyze corticostriatal synapses** A) Schematic illustration of IMARIS-based workflow to quantify corticostriatal synaptic density and overall morphology: Combined immunofluorescence (step 1) for VGLUT1 (excitatory, cortical terminal marker) and Homer1 (excitatory, postsynaptic marker) to stain for corticostriatal synapses in the striatum was followed by expansion microscopy (step 2) and confocal image acquisition (step 3). Generated confocal images were then 3D-rendered in IMARIS (step 4), followed by surface reconstruction (step 5). Synaptic loci were filtered as pre-and postsynaptic surfaces in close proximity to each other (<0.01 µm distance between surfaces). B) Histogram displaying pre- and postsynaptic shortest distances with peak indicating synaptic loci with shortest distance <0.01 µm.

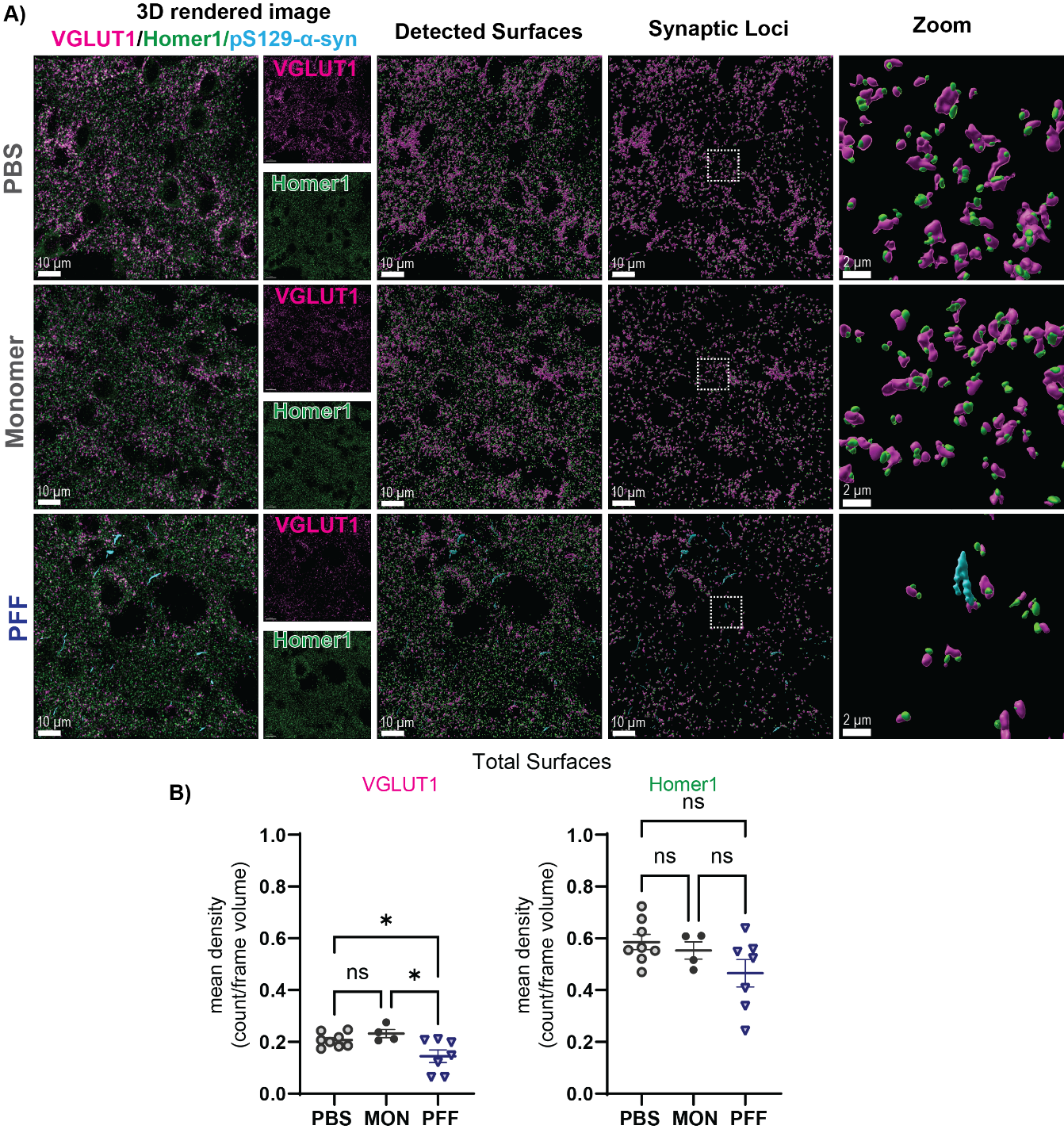
**Supplementary Figure 4: Traditional immunofluorescence protocol reveals reduction of corticostriatal VGLUT1/Homer1 synaptic loci in PFF injected mice** A) From left to right: Left panel showing 3D-rendered confocal images of VGLUT1 (magenta), Homer1 (green) and pS129-α-syn (cyan) for control (PBS, MON) and PFF-injected animals. The next panel shows detected surfaces for the markers of interest. Panels on the right show filtered synaptic loci consisting of VGLUT1/Homer1 surfaces in close proximity with inset to show juxtaposed positioning of corticostriatal synaptic markers. B) Quantification of density of total VGLUT1 and Homer1 surfaces: Total density of VGLUT1 surfaces was significantly reduced in PFF animals compared to control groups (1-way ANOVA test: F(2, 16)= 5.77, p=0.0227; Tukey‘s multiple comparison post hoc test: PBS vs PFF: p=0.0439, MON vs PFF: p=0.0187), while no changes in total Homer1 density were found between groups (1-way ANOVA test: F(2, 16)= 2.484, p=0.115). Statistical significance defined as: *p<0.05, **p<0.01, ***p<0.001, ****p<0.0001***. Corresponding statistical analysis information and group mean∓SEM are provided in table S1.

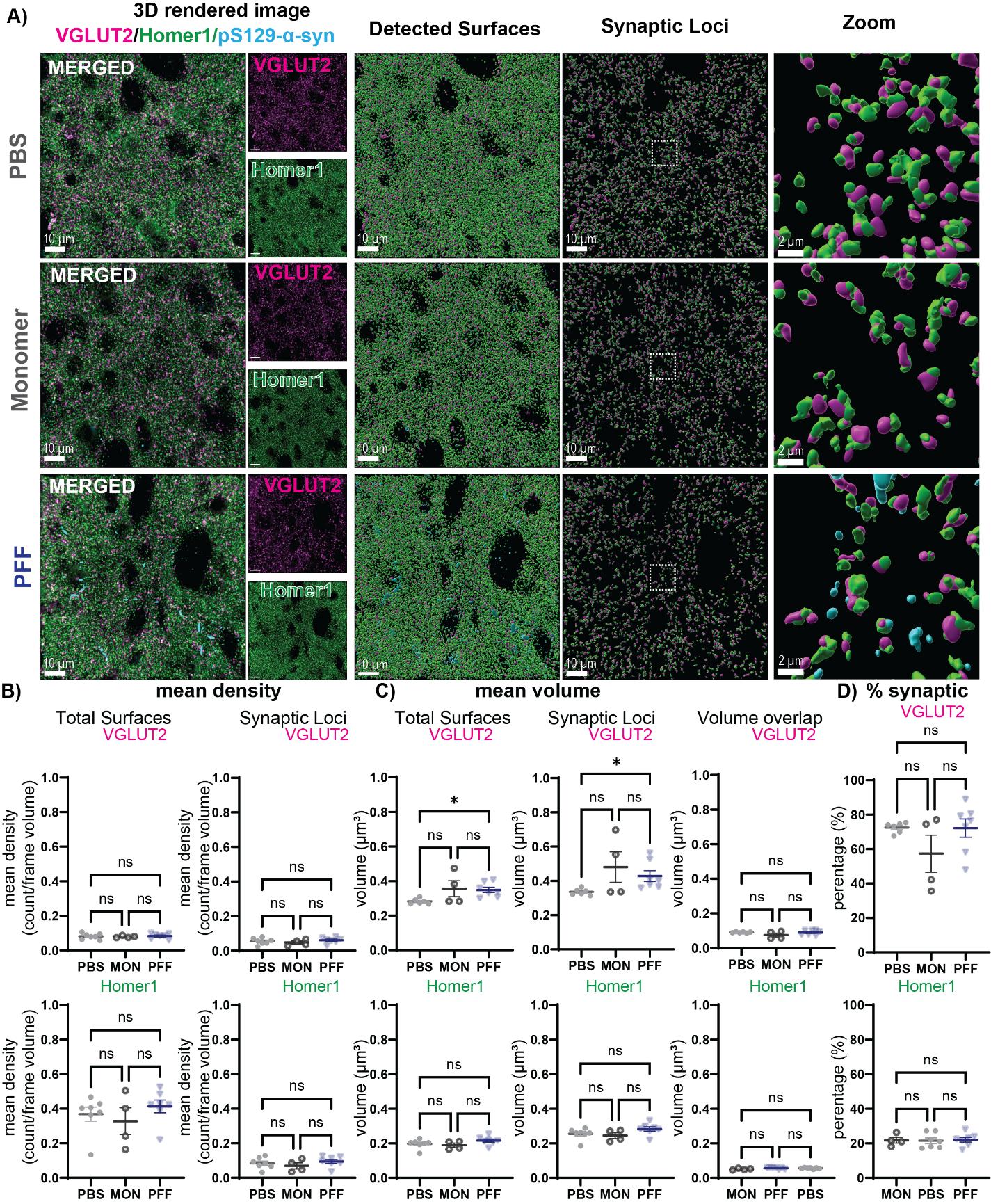
**Supplementary Figure 5: Striatal PFF injections affect morphology of thalamic inputs, but not density in the striatum** A) From left to right: Left panel showing 3D-rendered confocal images of thalamic terminal marker VGLUT2 (magenta), Homer1 (green) and pS129-α-syn (cyan) for control (Monomer, PBS) and PFF-injected animals. The next panel shows detected surfaces using IMARIS surface function for the markers of interest. Panels on the right show filtered synaptic loci consisting of close proximity VGLUT2/Homer1 surfaces with zoomed in frame to show juxtaposed positioning of thalamostriatal synaptic markers. D) Analysis of mean density of total surfaces and synaptic surfaces did not identify significant differences between groups (see table S1 for statistical overview). E) Quantification of thalamostriatal synaptic surface volumes: volumes for total VGLUT2 and synaptic VGLUT2 were significantly larger in PFF injected animals compared to PBS controls (total VGLUT2 volume: Welch’s ANOVA test: W(2, 5.917)= 7.673, p=0.0227; Dunnett‘s multiple comparison post hoc test: PBS vs PFF: p=0.0129; synaptic VGLUT2 volume: Kruskal Wallis test: p=0.0235; Dunnett‘s multiple comparison post hoc test: PBS vs PFF: p=0.0317), while neither total Homer1 volumes nor synaptic loci volumes were significantly different between groups (See table S1 for statistical overview). Volume overlap with juxtaposed surfaces was not changed between control groups and PFF animals. F) Percentage of synaptic loci for total surfaces did not differ between experimental groups. Statistical significance defined as: *p<0.05, **p<0.01, ***p<0.001, ****p<0.0001***. Corresponding statistical analysis information and group mean∓SEM are provided in table S2.

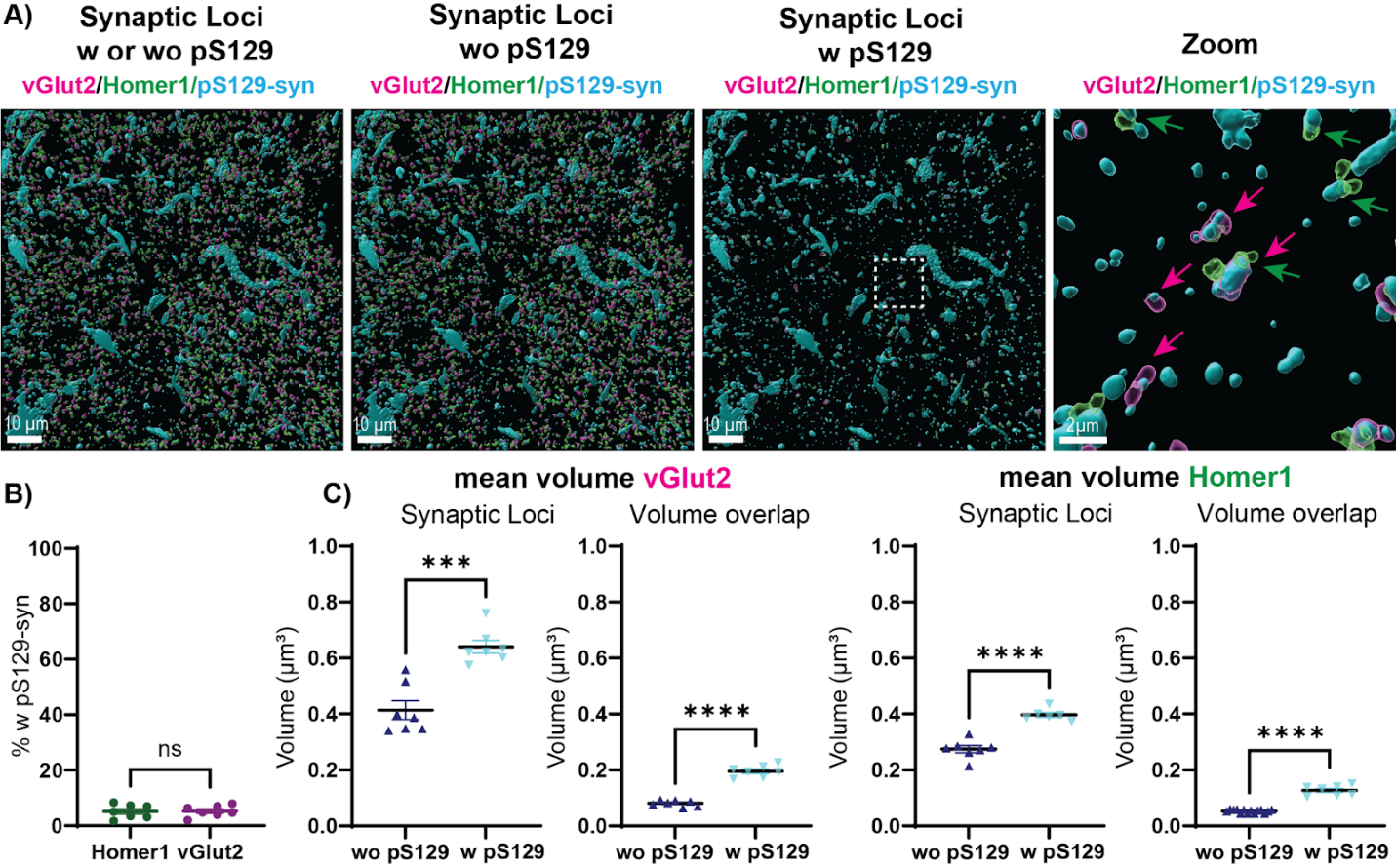
**Supplementary Figure 6: Morphological changes to pS129-α-syn-positive thalamostriatal synaptic compartments:** A) Shown are reconstructed, filtered corticostriatal synaptic loci (VGLUT2-Homer1) and pS129-α-syn pathology in the dorsal striatum of animals 6-weeks post PFF injections. Subsets of VGLUT2 and Homer1 synaptic loci showed co-localization with pS129-α-syn aggregates and are highlighted as transparent surfaces. Additional filtering allowed for the display of synaptic loci without pS129-α-syn (wo pS129) aggregates and synaptic loci with pS129-α-syn (w pS129) aggregates. A zoom-in of synaptic loci w pS129 highlights synaptic VGLUT2 w pS129 (magenta arrow and surface) and synaptic Homer1 w pS129 (green arrow and surface), pathologic pS129-α-syn aggregates are shown in cyan. B) Quantification of percentage of synaptic loci with pS129-α-syn aggregates: no difference in positive percentage for small, intrasynaptic pS129-α-syn aggregates between VGLUT2 and Homer1 synaptic loci. (Unpaired Student’s t-test, p=0.9334). C) Volumes of synaptic loci positive (w pS129) for pS129-α-syn-aggregates show a significant enlargement in volumes for VGLUT2 (Unpaired Student’s t-test, p=0.0001) and Homer1(Unpaired Student’s t-test, p<0.0001). Volume overlapped with respective synaptic partners of synaptic loci w pS129 were significantly larger compared to synaptic loci wo pS129 (VGLUT2: Unpaired Student’s t-test, p<0.0001, Homer1: Welch t-test, p<0.0001). The percentage of synaptic loci for total surfaces did not differ between experimental groups. Statistical significance defined as: *p<0.05, **p<0.01, ***p<0.001, ****p<0.0001***. Corresponding statistical analysis information and group mean∓SEM are provided in table S3.

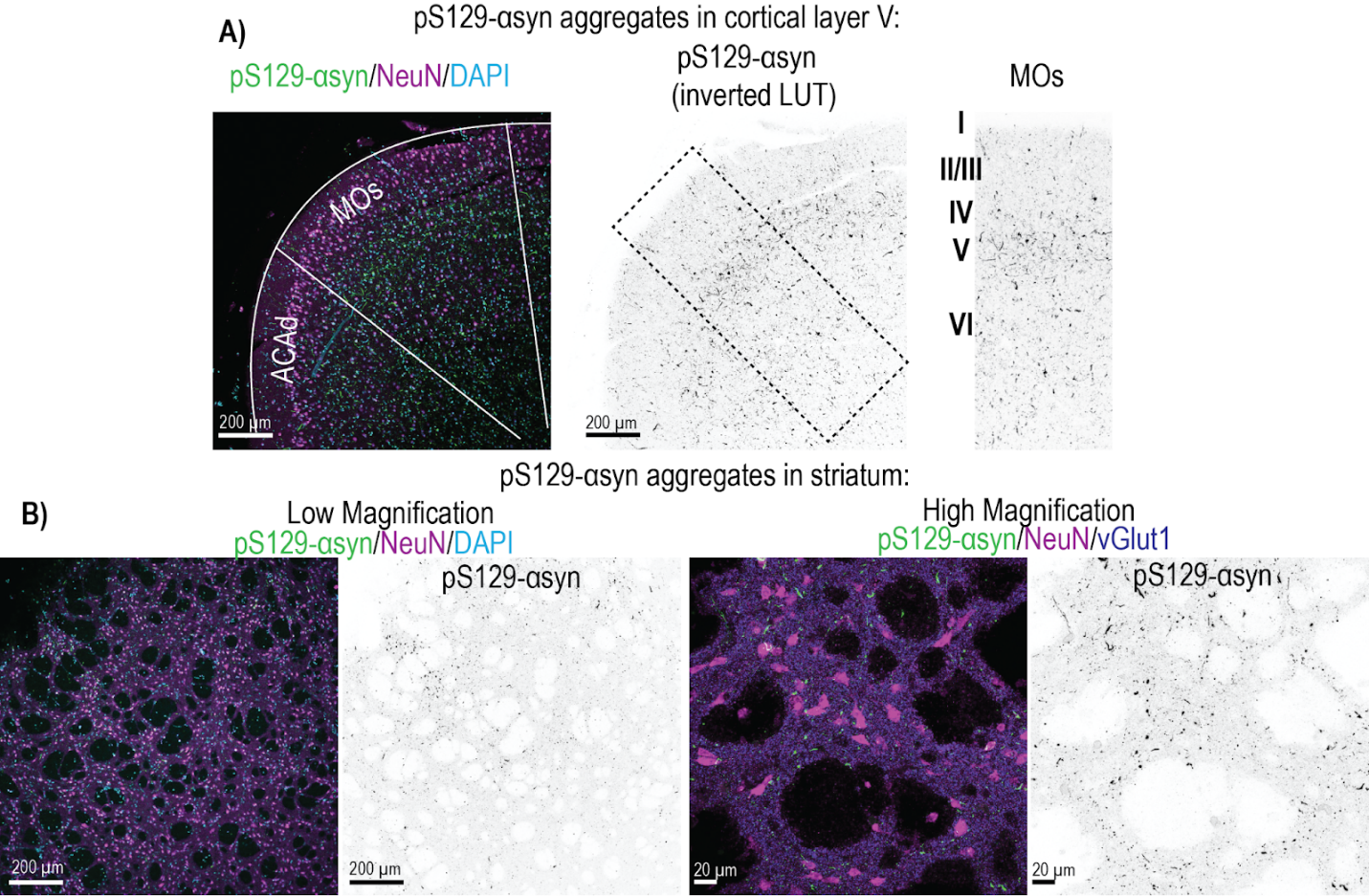
**Supplementary Figure 7: Posthoc pathology staining in representative PFF-treated brain slice from physiological recordings** A) Representative immunofluorescence microscopy images of the MOs and ACAd in a slice post fixated after electrophysiological recordings to qualitatively assess pathology formation in hypothesis-relevant brain areas. A distinct topographical accumulation of pS129-ɑ-syn positive signal (green) is visible right after the NeuN-dense (magenta) cortical layer 2/3 (left side). LUT-inverted image (middle) of pS129-ɑ-syn signal highlights the specific accumulation of ɑ-syn aggregates, which is concentrated to deeper cortical layers. Scale bar: 200 µm. The right side shows a zoom-in of the MOs with cortical layer annotation, highlighting the distinct pathology accumulation in layer 5 and layer 6. B) Low magnification (left) and high magnification (right) confocal images showing abundant pS129-ɑ-syn pathology (green) in the dorsal striatum of post-hoc stained section from physiological recordings.

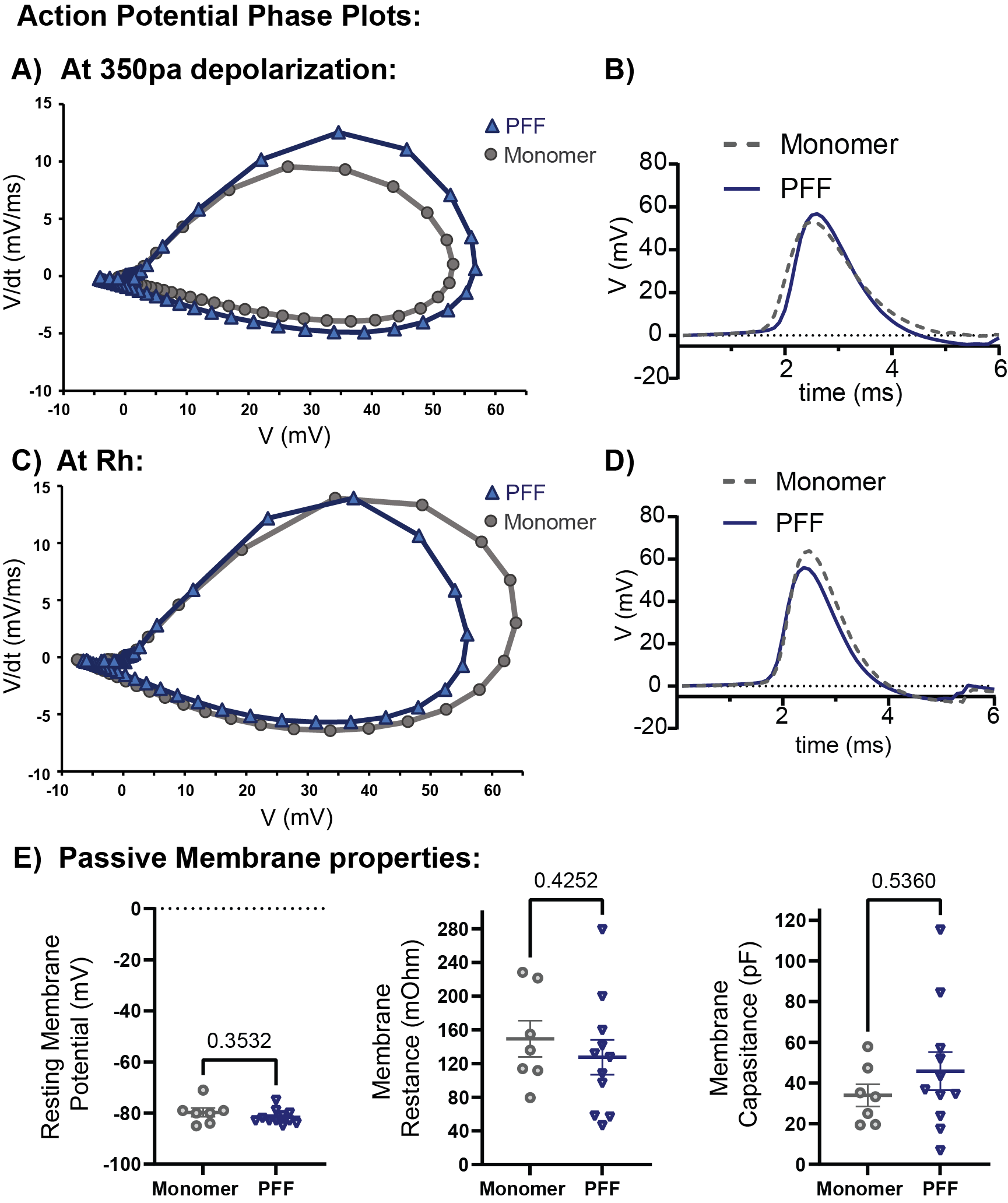

**Supplementary Figure 8: PFF injections do not change in intrinsic membrane properties**

**Table S4** A) Shown are AP phase plots from whole cell recordings of SPNs in monomer (gray) or PFF (blue) injected groups, and B) shows the average AP shape for the different groups at 350pA depolarizing stimulation. C+D) Shown AP phase plots and average AP shape for different treatment at Rh current injection. E) Passive membrane properties did not change between control and PFF injected group (Student’s t-test). Statistical significance defined as: *p<0.05, **p<0.01, ***p<0.001, ****p<0.0001***. Corresponding statistical analysis information and group mean∓SEM are provided in table S4.

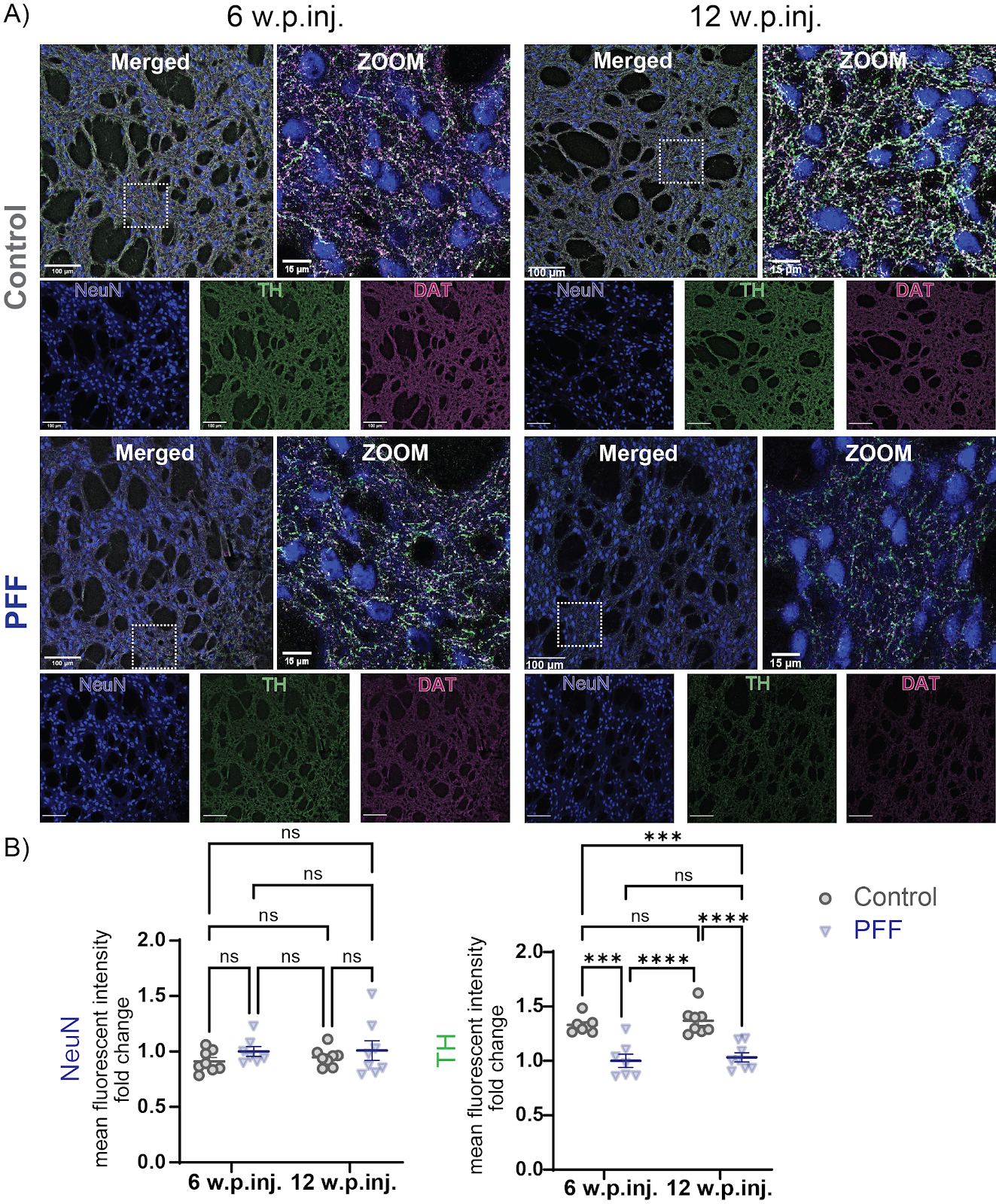
**Supplementary Figure 9 Early dopaminergic denervation of the dorsal striatum in PFF injected mice.** A) Representative immunofluorescence microscopy images of the dorsal striatum for 6-week (left side) and 12 week (right side) striatally PFF- (lower panel) and control (PBS)-injected animals stained for the dopaminergic terminal markers tyrosine hydroxylase (TH, green) and dopamine transporter (DAT, magenta) and neuronal marker NeuN (blue) as normalization control. Scale bars: 100 µm and 15 µm for zoomed in frames. B) Quantification of mean fluorescent intensities of markers of interest: mean fluorescent intensity of NeuN did not change between treatment groups nor for time points (Ordinary 2-way ANOVA). TH signal was normalized to NeuN mean fluorescent intensity and shown as fold change of the 6-week PFF group. Normalized TH mean fluorescent intensity fold change was significantly reduced in PFF injected animals compared to control at the 6-weeks and 12 weeks post injection timepoints. No main effect of time point post injection was observed for TH fluorescent intensity for the different treatment groups. (Ordinary 2-way ANOVA, Column Factor: F(1, 27)=55.35, p<0.0001, Tukey’s multiple comparison post hoc test). Statistical significance defined as: *p<0.05, **p<0.01, ***p<0.001, ****p<0.0001***. Corresponding statistical analysis information and group mean∓SEM are provided in table S5.

| **Test** | | **[group] (Mean±SEM), N** | **F** | **Dfn** | **Dfd** | **p** |
| --- | --- | --- | --- | --- | --- | --- |
| Total VGLUT1 Density | Ordinary one-way ANOVA | PBS (0.2068±0.0.0097), N=8; Monomer (0.2322±0.0159), N=4; PFF (0.1447±0.0.0242), N=7; | 5.77 | 2 | 16 | 0.013 |
|  | Tukey's multiple comparison post hoc test | PBS vs PFF |  |  |  | **0.0439** |
|  | Tukey's multiple comparison post hoc test | Monomer vs PFF |  |  |  | **0.0187** |
| Total Homer1 Density | Ordinary one-way ANOVA | PBS (0.5856±0.0.0296), N=8; Monomer (0.5530±0.0331), N=4; PFF (0.4653±0.0529), N=7; | 2.484 | 2 | 16 | 0.115 |

**Table S1: Statistics Summary Table for Figure S4**

|  | Test | [group] (Mean±SEM), N | F | Dfn | Dfd | p |
| --- | --- | --- | --- | --- | --- | --- |
| Total VGLUT2 Density | Ordinary one-way ANOVA | PBS (0.081±0.001), N=7; Monomer (0.079±0.003), N=4; PFF (0.083±0.006), N=7 | 0.08003 | 2 | 15 | 0.9235 |
| Total Homer1 Density | Welch's ANOVA test | PBS (0.408±0.015), N=6*; Monomer (0.328±0.077), N=4; PFF (0.414±0.036), N=7 | W=0.4875 | 2 | 6.036 | 0.6364 |
| Synaptic VGLUT2 Density | Ordinary one-way ANOVA | PBS (0.054±0.007), N=7; Monomer (0.058±0.008), N=4; PFF (0.060±0.007), N=7 | 0.2025 | 2 | 15 | 0.8189 |
| Synaptic Homer1 Density | Ordinary one-way ANOVA | PBS (0.084±0.012), N=7; Monomer (0.070±0.018), N=4; PFF (0.095±0.012), N=7 | 0.48 | 2 | 15 | 0.7711 |
| Total VGLUT2 Volume | Welch's ANOVA test | PBS (0.28±0.01), N=7; Monomer (0.36±0.05), N=4; PFF (0.035±0.02), N=7 | W=7.673 | 2 | 5.917 | 0.0227 |
|  | Dunnett's multiple comparison post hoc test | PBS vs PFF |  |  |  | 0.0129 |
| Total Homer1 Volume | Ordinary one-way ANOVA | PBS (0.20±0.01), N=7; Monomer (0.19±0.01), N=4; PFF (0.21±0.01), N=7 | 1.704 | 2 | 15 | 0.2154 |
| Synaptic VGLUT2 Volume | Kruskal-Wallis test | PBS (0.34±0.01), N=6*; Monomer (0.48±0.09), N=4; PFF (0.43±0.03), N=7 |  |  |  | 0.0235 |
|  | Dunnett's multiple comparison post hoc test | PBS vs PFF |  |  |  | 0.0317 |
| Synaptic Homer1 Volume | Kruskal-Wallis test | PBS (0.25±0.01), N=7; Monomer (0.25±0.02), N=4; PFF (0.28±0.01), N=7 |  |  |  | 0.872 |
| Synaptic VGLUT2 Volume Overlapped | Welch's ANOVA test | PBS (0.090±0.001), N=6*; Monomer (0.075±0.010), N=4; PFF (0.089±0.005), N=7 | W=0.9232 | 2 | 5.724 | 0.3173 |
| Synaptic Homer1 Volume Overlapped | Ordinary one-way ANOVA | PBS (0.0560±0.001), N=7; Monomer (0.051±0.002), N=4; PFF (0.056±0.002), N=7 | 2.28 | 2 | 15 | 0.1366 |
| % Synaptic VGLUT2 | Welch's ANOVA test | PBS (72.49±1.25), N=6*; Monomer (57.34±10.76), N=4; PFF (72.16±5.35), N=7 | W=0.8745 | 2 | 5.179 | 0.4106 |
| % Synaptic Homer1 | Ordinary one-way ANOVA | PBS (21.50±1.63), N=6*; Monomer (21.75±1.69), N=4; PFF (22.10±1.40), N=7 | 0.04136 | 2 | 15 | 0.9596 |

**Table S2: Statistics Summary Table for Figure S5** *: Outlier removed (Grout test)

|  | Test | [group] (Mean±SEM), N | t | df | p |
| --- | --- | --- | --- | --- | --- |
| %w pS129 in Homer1 vs VGLUT2 | Unpaired Student's t-test | Homer1 (5.13±0.94), N=7; VGLUT2 (5.24±0.77), N=7 | 0.08535 | 12 | 0.9334 |
| Synaptic VGLUT2 Volume | Unpaired Student's t-test | wo pS129 (0.41±0.03), N=7; w pS129 (0.64±0.02), N=7 | 3.807 | 12 | 0.0001 |
| Synaptic VGLUT2 Volume Overlapped | Unpaired Student's t-test | wo pS129 (0.081±0.004), N=7; w pS129 (0.196±0.008), N=7 | 13.08 | 12 | <0.0001 |
| Synaptic Homer1 Volume | Unpaired Student's t-test | wo pS129 (0.27±0.02), N=7; w pS129 (0.40±0.01), N=6* | 5.013 | 11 | <0.0001 |
| Synaptic Homer1 Volume Overlapped | Unpaired Student's t-test, Welch Correction | wo pS129 (0.053±0.002), N=7; w pS129 (0.127±0.007), N=7 | 10.39 | 7.424 | <0.0001 |

**Table S3: Statistics Summary Table for Figure S6** *: Outlier removed (Grout test)

| Test |  | [group] (Mean±SEM), N | t | df | p |
| --- | --- | --- | --- | --- | --- |
| Resting Membrane Potential | Unpaired Student's t-test | Monomer (-79.71±1.71), n=7, N=3; PFF (-81.64±0.85), n=11, N=5 | 1.119 | 16 | 0.2797 |
| Membrane Resistance | Unpaired Student's t-test | Monomer (149.4±21.4), n=7, N=3; PFF (127±20.7), n=11, N=5 | 0.7019 | 16 | 0.4929 |
| Membrane Capacitance | Unpaired Student's t-test | Monomer (33.94±5.46), n=7, N=3; PFF (45.83±9.37), n=11, N=5 | 0.9424 | 16 | 0.36 |

**Table S4: Statistics Summary Table for Figure S8**

|  | Test | [group] (Mean±SEM), N | F | Dfn | Dfd | p |
| --- | --- | --- | --- | --- | --- | --- |
| NeuN Mean Fluorescent Intensity | Ordinary two-way ANOVA | 6wk PBS (0.911±0.034), N=8; 6wk PFF (1.000±0.044), N=7; 12wk PBS (0.948±0.030), N=8; 12wk PFF (1.009±0.089), N=8; Row Factor (Time Point)  Column Factor (Treatment) Interaction | 0.1792 1.853 0.06508 | 1  1  1 | 27 27 27 | 0.6755 0.1847 0.8006 |
| TH Mean Fluorescent Intensity (norm. to NeuN) | Ordinary two-way ANOVA | 6wk PBS (1.366±0.043), N=8; 6wk PFF (1.000±0.060), N=7; 12wk PBS (1.369±0.044), N=8; 12wk PFF (1.033±0.042), N=8; Row Factor (Time Point)  Column Factor (Treatment) Interaction | 0.1477 55.35 0.1026 | 1  1  1 | 27 27 27 | 0.7038 <0.0001 0.7513 |
| 6wk PBS vs 6wk PFF | Tukey's multiple comparison post hoc test | 6wk PBS (1.366±0.043), N=8; 6wk PFF (1.000±0.060), N=7 |  |  |  | <0.0001 |
| 6wk PBS vs 12wk PFF | Tukey's multiple comparison post hoc test | 6wk PBS (1.366±0.043), N=8; 12wk PFF (1.033±0.042), N=8 |  |  |  | 0.0001 |
| 6wk PFF vs 12wk PBS | Tukey's multiple comparison post hoc test | 6wk PFF (1.000±0.060), N=7; 12wk PBS (0.948±0.030), N=8 |  |  |  | <0.0001 |
| 12wk PBS vs 12wk PFF | Tukey's multiple comparison post hoc test | 12wk PBS (0.948±0.030), N=8; 12wk PFF (1.033±0.042), N=8 |  |  |  | 0.0001 |

**Table S5: Statistics Summary Table for Figure S9**
